# Compensatory CSF2-driven macrophage activation promotes adaptive resistance to CSF1R inhibition in breast-to-brain metastasis

**DOI:** 10.1101/2021.06.07.447034

**Authors:** Florian Klemm, Alexander Schäffer, Anna Salamero-Boix, Tijna Alekseeva, Michael Schulz, Katja Niesel, Roeltje R. Maas, Marie Groth, Benelita T. Elie, Robert L. Bowman, Monika E. Hegi, Roy T. Daniel, Pia S. Zeiner, Jenny Zinke, Patrick N. Harter, Karl H. Plate, Johanna A. Joyce, Lisa Sevenich

## Abstract

Tumor microenvironment-targeted therapies are emerging as promising treatment options for different cancer types. Tumor-associated macrophages and microglia (TAMs) represent an abundant non-malignant cell type in brain metastases and have been proposed to modulate metastatic colonization and outgrowth. We used an inhibitor of colony stimulating factor 1 receptor (CSF1R) to target TAMs at distinct stages of the metastatic cascade in preclinical breast-to-brain metastasis models and found that CSF1R inhibition leads to anti-tumor responses in prevention and intervention trials. However, in established brain metastases, compensatory CSF2Rb-STAT5-mediated pro-inflammatory TAM activation blunted the ultimate efficacy of CSF1R inhibition by inducing neuro-inflammation gene signatures in association with wound repair responses that fostered tumor recurrence. Consequently, combined blockade of CSF1R and STAT5 signaling led to sustained tumor control, a normalization of microglial activation states and amelioration of neuronal damage.

## INTRODUCTION

Brain metastases (BrM) represent the most common intracranial tumors in adults. It is estimated that approximately 20-40% of all cancer patients develop BrM, with melanoma, breast, and lung cancer showing the highest incidence ^1–4^. Despite recent advances in the management of extracranial primary tumors, treatment options for disseminated disease remain limited and largely depend on multimodal approaches involving surgical resection, radio- and/or chemotherapy ^5,6^. The success of immunotherapies in several non-cerebral cancers has led to considerable optimism for the development of more efficient treatment options for brain tumors, including glioblastoma and BrM ^7,8^. However, strategies that aim to activate T cell responses directed against tumor cells have largely failed as monotherapies in primary brain tumors and showed low or moderate efficacy in BrM patients depending on the primary tumor type ^9–11^. Low response rates can in part be attributed to the unique tissue environment of the central nervous system (CNS), which is highly immune-suppressive ^12,13^.

Tumor-associated macrophages (TAMs) are one of the most abundant non-cancerous cell types in breast-BrM ^14–16^ and originate from brain-resident microglia (TAM-MG) or monocyte-derived macrophages (TAM-MDM) ^14,15,17,18^ which exhibit an immunomodulatory phenotype in BrM ^14^. Therefore, targeting this highly abundant TAM population could represent a viable strategy to disrupt cancer-permissive tumor-stroma interactions and to modulate the immune response directed against cancer cells for more effective tumor control. Indeed, preclinical studies have demonstrated that TAM-targeted therapies can lead to anti-tumor efficacy in glioblastoma ^19–21^, melanoma-BrM ^22^, and in the context of intracerebral injection of 4T1 breast cancer cells ^23^. However, these previous studies interrogated the role of TAMs in BrM by depleting the cells only at single time points in the metastatic process and did not discriminate cell-type specific functions or the consequences of CSF1R blockade on TAM-MG and TAM-MDM, which may have important implications for clinical translation. Thus, in order to address this knowledge gap, we used an inhibitor of colony stimulating factor 1 receptor (CSF1R) to deplete TAMs at distinct stages of BrM, to study their role in metastatic progression, and to evaluate the efficacy of TAM-targeted therapies for BrM in different preclinical trial settings.

## RESULTS

### CSF1R expression in human and mouse brain metastases

We first analyzed the abundance of CSF1R+ cells in patient BrM tissue arrays originating from different cancer types including small cell lung cancer (SCLC), non-small cell lung cancer (NSCLC), renal cell cancer (RCC), melanoma, colon and breast cancer (Extended data Table 1). We observed high infiltration of CSF1R+ immune cells across all analyzed BrM lesions, with RCC-BrM showing the largest proportion of CSF1R+ cells, and SCLC-BrM the lowest (Fig. 1a,b). Localization of CSF1R+ cells showed a considerable overlap with IBA1 expression in all analyzed biopsies, indicating that the majority of CSF1R+ cells originate from the monocyte/macrophage lineage. Further investigation in an independent cohort of breast cancer-BrM and NSCLC-BrM whole tissue sections (Fig. 1c, Extended data Fig. 1, Extended data Table 2) confirmed CSF1R expression in a significant proportion of BrM-MG and BrM-MDMs, whereas very few non-immune cells were CSF1R+. To examine the potential relevance of low level CSF1R expression in tumor cells, we quantified *CSF1R/Csf1r* across a panel of human and mouse cancer cell lines. Quantitative real-time PCR showed either no or very low *CSF1R/Csf1r* expression in cancer cell lines derived from different primary tumors, compared to macrophages as a positive control (Extended data Fig. 2a,b). Moreover, *CSF1R* expression levels in FACS-purified tumor cells isolated from experimental BrM lesions were significantly lower than from tumor-associated MG and MDM or their normal counterparts (Extended data Fig. 2c). Finally, tumor cell viability was not affected by treatment with the CSF1R inhibitor BLZ945 *in vitro* (Extended data Fig. 2d), indicating that even though rare tumor cells can express low levels of CSF1R, their survival is not directly dependent on CSF1R signaling, and its inhibition rather targets myeloid cells.

**Fig. 1.**
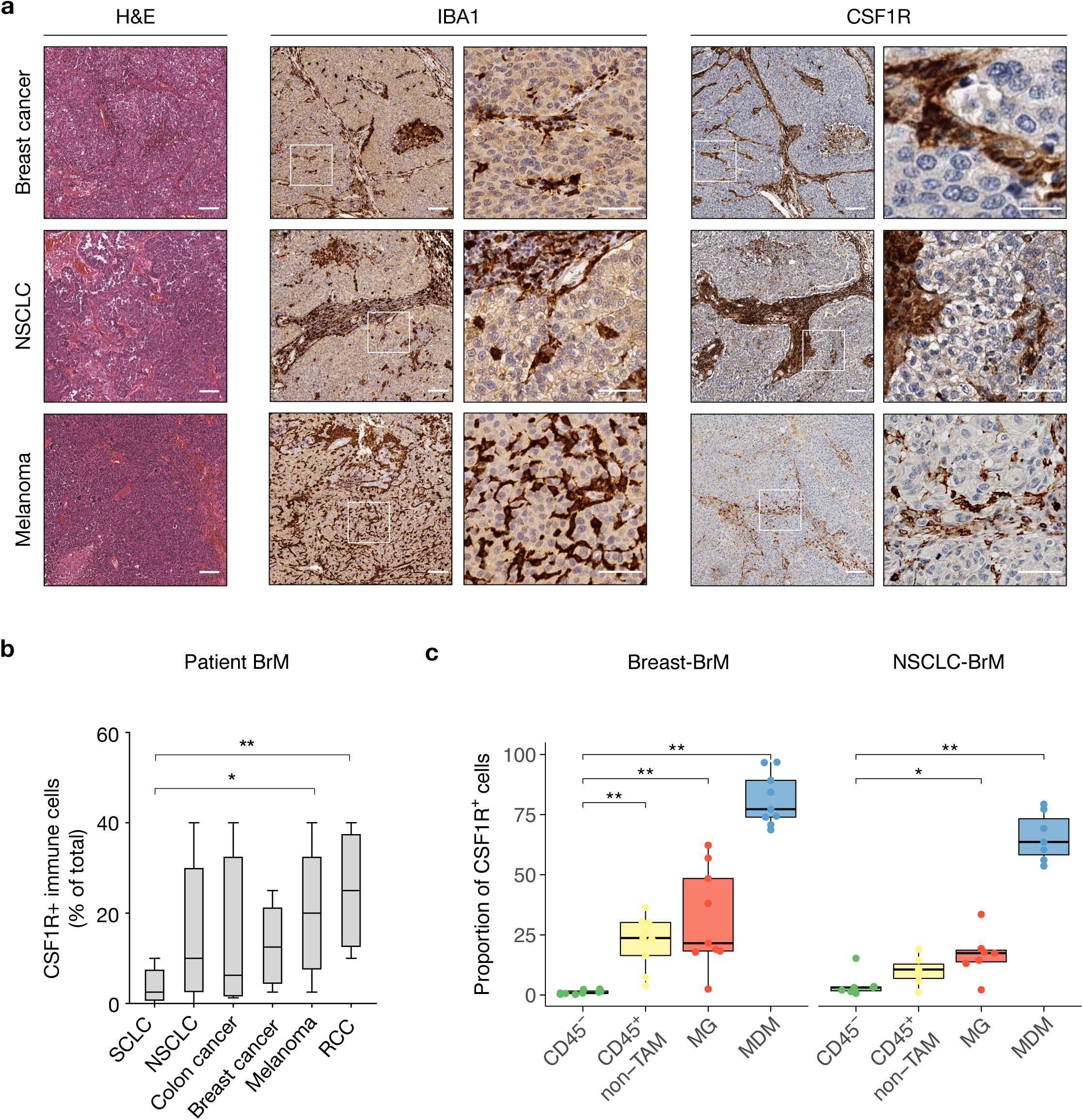
Tumor-associated microglia and macrophages (TAMs) in human BrM express CSF1R. **a,** Representative IHC images of IBA1+ and CSF1R+ cells in human brain metastasis (BrM) samples. Scale bar, 100 and 50 μm. **b,** Quantification of CSF1R+ immune cells in BrM originating from NSCLC (n=43), SCLC (n=9), colon (n=4), breast (n=26), melanoma (n=14), and renal cancer (n=5). **c,** Percentage of CSF1R+ cells in CD45^-^ non-immune cells (CD45^-^), CD45^+^ non-TAM immune cells (CD45^+^, CD68/P2RY12^-^, CD49D^-/+^), MG (CD45^+^, CD68/P2RY12^+^, CD49D^-^) and MDM (CD45^+^, CD68/P2RY12^+^, CD49D^+^) in BrM originating from breast (n=9) and NSCLC (n=7), as determined by immunofluorescence (IF). Scale bar, 100 μm. \**P*<0.05, \*\**P*<0.01; two-tailed Student’s t-test in (b), one-sided Wilcoxon signed-rank test in (c).

### Ontogenetic origin and molecular identity of TAMs in BrM

We next analyzed the localization and morphology of TAMs in the xenograft MDA-BrM and syngeneic 99LN-BrM2 breast-BrM mouse models ^24^. TAM activation was associated with a gradual change in cell morphology from ramified, resting MG in the adjacent brain parenchyma to amoeboid, activated TAMs within the tumor lesion (Fig. 2a). Flow cytometric analysis using previously established lineage-specific cell surface markers ^17^, revealed that the majority of TAMs in BrM were derived from brain-resident MG, while on average 7% and 17% were recruited from the periphery in the MDA-BrM and 99LN-BrM models respectively (Extended data Fig. 2e). We performed RNA sequencing (RNAseq) analyses of FACS-purified TAM-MG and TAM-MDM from MDA-BrM tumors compared to their respective control population in tumor-free animals, i.e. normal MG or blood monocytes, to gain deeper insight into the molecular identity of TAMs in BrM (Fig. 2b,c, see also Extended data Table S3). The purity of the FACS-sorted populations was further validated by the MG- and MDM-specific expression of cell type-restricted markers (Extended data Fig. 2f). RNAseq analysis revealed 966 differentially expressed genes (DEG) in the TAM-MG population compared to normal MG, and 2863 DEG for the comparison between TAM-MDM and normal monocytes; 500 of which were shared between both TAM populations (Fig. 2b and Extended data Table 3). We also queried an RNAseq dataset from matched primary breast cancer and BrM bulk tissue samples ^25^ for the expression of the 3329 DEG identified in the comparison of TAMs to normal MG and monocytes in the mouse model. Interestingly, we found a higher enrichment of the metastasis TAM genes in BrM compared to the matched primary breast cancer both in triple negative breast cancer (TNBC) and in other molecular subtypes collectively denoted as non-TNBC (Extended data Fig. 2g). This suggests that these TAM-DEG are acquired in a breast BrM-specific fashion. We next performed gene ontology analysis of clustered gene groups within the 500 genes commonly enriched in both TAM populations (Fig. 2c and Extended data Table 4). TAM-MDM showed a higher expression of gene signatures that represent pathways associated with antigen presentation, and tissue remodeling and immune regulation (cluster 5, 6 and 8) as well as mitosis and cell division (cluster 7), while TAM-MG showed a higher enrichment of genes associated with wounding responses and regulation of neurological processes (cluster 3). Signatures associated with negative regulation of CNS development, metabolism and leukocyte recruitment (cluster 1, 2 and 4) showed similar induction in TAM-MG and TAM-MDM. Taken together, our data indicate that education of TAMs in BrM leads to the induction of cancer-associated inflammation that has previously been associated with tumor promotion across different cancer types ^26–29^. However, the functional contributions of TAM-MG and TAM-MDM at distinct stages of breast-to-brain metastasis remains largely unknown.

**Fig. 2.**
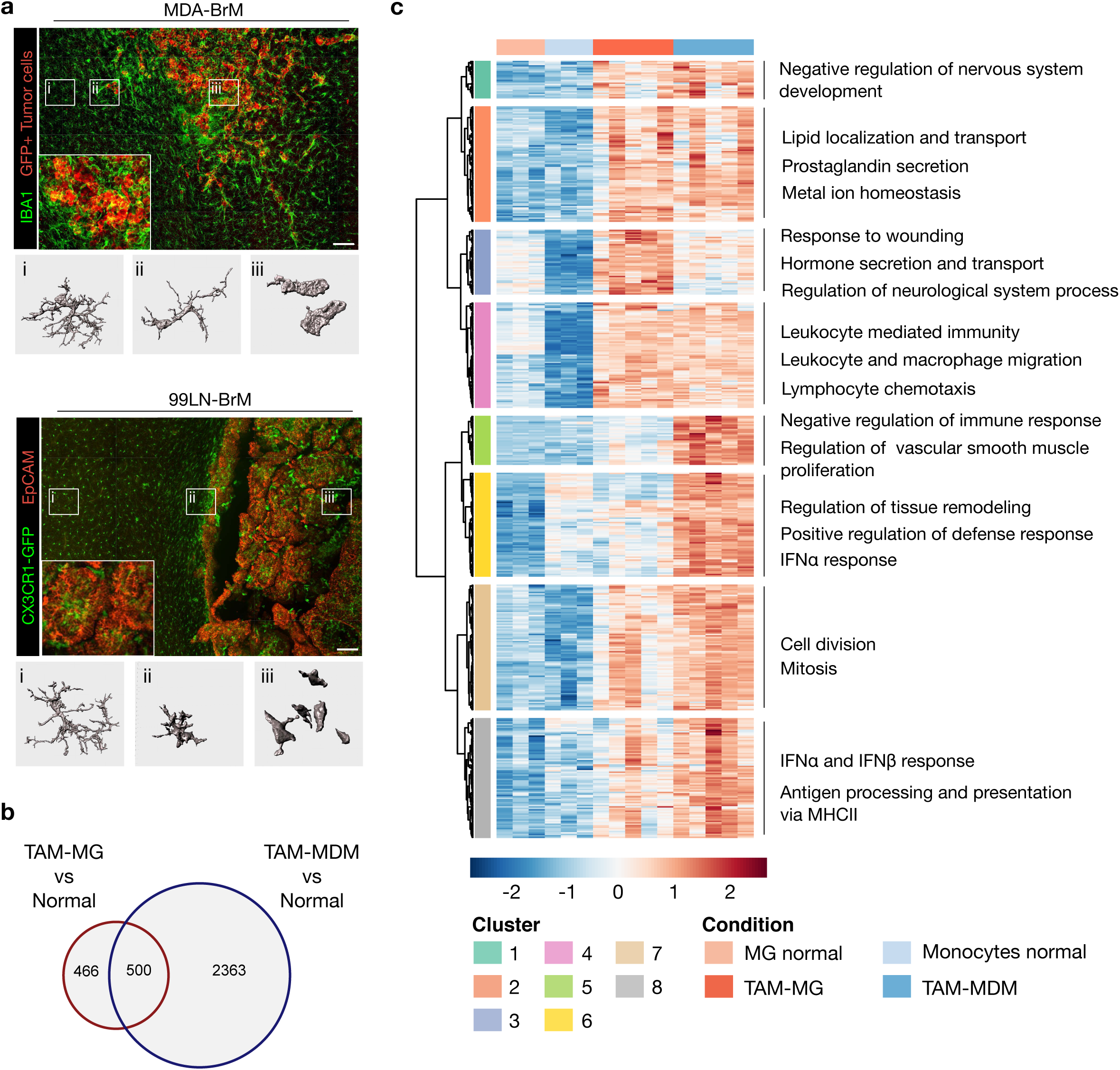
TAMs exhibit an inflammatory phenotype in mouse BrM. **a,** Representative IF images of TAMs in MDA-BrM (upper panel) and 99LN-BrM (lower panel), Scale bar, 250 μm. 3D image reconstructions of normal microglia (MG) and TAMs are shown in panels (i, ii, iii) below. **b,** Euler plot of the intersection of MG and monocyte-derived macrophage (MDM) genes upregulated in TAMs in the MDA-BrM model compared to normal controls (i.e. microglia and monocytes respectively from untreated, non-tumor-bearing mice). **c,** Gene expression heatmap of 500 upregulated genes in TAMs, compared to their normal cellular counterparts. Rows were clustered using Ward’s method with 1 – Pearsońs correlation coefficient as the distance measurement. Gene ontology terms found overrepresented by gene set overrepresentation analysis (ORA) within each cluster are indicated.

### MG support tumor cell extravasation

We next sought to investigate the consequences for tumor cell fate after initial contact with MG. We therefore employed a MG depletion strategy using the blood-brain barrier (BBB)-permeable CSF1R inhibitor BLZ945 ^19^, and began by treating normal, non-tumor bearing mice which resulted in a 90% decrease in MG after 7 days (Fig. 3a,b). To dynamically track the first encounter of MG and tumor cells in the brain parenchyma, we performed *ex vivo* brain slice assays (Fig. 3c). MG in brain slices that were isolated from vehicle-treated mice displayed directed movements towards exogenously-added 99LN-BrM cells and extended their protrusions to survey the tumor cell surface (Fig. 3d and Extended data Movie 1), consistent with their known role in immune surveillance ^30^. The MG that remained in brain slices from BLZ945-treated animals were still motile and the formation of protrusions was visible. However, the execution of movements was less targeted, suggesting that following CSF1R inhibition MG failed to detect the presence of tumor cells (Fig. 3d and Extended data Movie 2). Indeed, quantification of tumor cell–MG interactions, normalized to the number of MG present following BLZ945 treatment, revealed a significant reduction in direct contacts (Fig. 3e). We also detected fewer tumor cells after 20h in culture in brain slices isolated from BLZ945-treated animals, compared to vehicle controls (Fig. 3f), suggesting that the initial tumor cell-MG interactions in the controls result in the support of tumor cell colonization rather than recognition and killing of tumor cells.

**Fig. 3.**
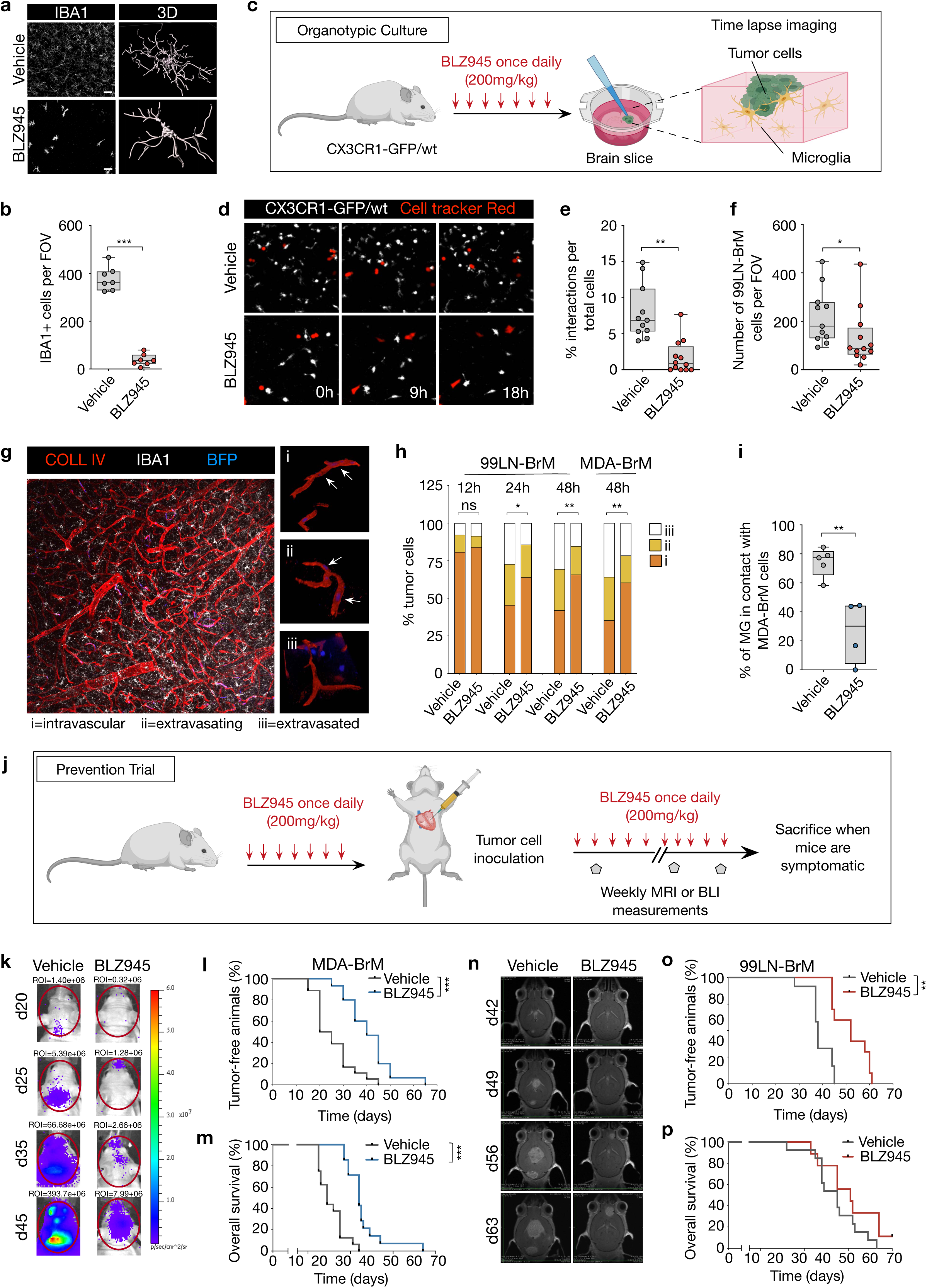
MG support initial steps of metastatic colonization. **a,** Representative IF images (left) and 3D reconstruction (right) of IBA1+ microglia in normal brain parenchyma treated for 7 days with vehicle or BLZ945. Scale bar, 25 μm. **b,** Quantification of IBA1+ cells after vehicle or BLZ945 treatment for 7 days, n=7 per group with 5 analyzed fields of view (FOV) acquired with a 10x objective. **c,** Experimental design for time-lapse imaging of MG–tumor cell interactions on brain slices isolated from vehicle or BLZ945-treated mice. **d,** Representative images of MG-tumor cell interactions on brain slices at different time points during time-lapse imaging. (n=3 independent experiments). **e,** Quantification of the number of interactions between tumor cells and MG (n=11; vehicle, n=12; BLZ945), normalized to the total number of MG in each condition. **f,** Quantification of tumor cells on brain slices after 20h (n=11; vehicle, n=12; BLZ945). **g,** Representative image of a brain section stained for collagen IV (COLL IV, red) and IBA1 (white) to visualize blood vessels and microglia respectively. Insets on right depict the relative localization of tumor cells (BFP, blue) to the vasculature. Arrows indicate intravascular and extravasating cells. **h,** Quantification of tumor cell extravasation in response to BLZ945 treatment at 12h, 24h and 48h after tumor cell injection (n=3-5 mice per condition). **i,** Quantification of direct contacts between MDA-BrM tumor cells and MG at sites of extravasation (n=5; vehicle, n=4; BLZ945), normalized to the total number of MG. **j,** Experimental design of prevention trials. **k,** Representative BLI images of vehicle and BLZ945-treated MDA-BrM mice at different time points after tumor cell injection. **l,m,** Kaplan-Meier curves show the percentage of tumor-free animals **(l)** and the overall survival **(m)** in the MDA-BrM model (n=16; vehicle, n=14; BLZ945). **n,** Representative MRI images of vehicle and BLZ945-treated 99LN-BrM mice at different time points after tumor cell injection. **o,p,** Kaplan-Meier curves show the percentage of tumor-free animals **(o)** and the overall survival **(p)** in the 99LN-BrM model (n=12; vehicle, n=10; BLZ945). \**P*<0.05, \*\**P*<0.01, \*\*\**P*<0.001; two-tailed Student’s t-test in (b, e, f, i), Chi square test in (h), Mantel-Cox log rank test in (l, m, o, p).

To further evaluate if these tumor cell-MG interactions are also apparent at the BBB *in vivo*, we treated mice with BLZ945 or vehicle for 7 days before intracardiac injection of tumor cells and analyzed their rate of extravasation in the brain. We observed a significant delay in extravasation in BLZ945-treated 99LN-BrM animals after 24 and 48h (Fig. 3g,h). Similar results were observed in the MDA-BrM model after 48h (Fig. 3h). Consistent with the observations from the *ex vivo* brain slice assay with exogenously-added tumor cells, we detected fewer direct interactions between tumor cells and MG at sites of extravasation in BLZ945-treated mice, again normalized to the number of MG (Fig. 3i). *In vitro* BBB assays further confirmed that the addition of freshly isolated primary MG, or the MG cell line EOC2, increased the BBB transmigration potential of 99LN-BrM and MDA-BrM tumor cell lines (Extended data Fig. 3a-c), and this effect was significantly reduced by BLZ945 (Extended data Fig. 3b,c).

### TAMs support metastatic colonization and outgrowth

In order to evaluate whether the impaired BBB transmigration translates into decreased BrM incidence or delayed tumor outgrowth, we followed BrM progression in response to BLZ945 treatment. CSF1R inhibition in the prevention trial setting resulted in 94% and 73% reduction of the number of Iba+ cells in MDA-BrM and 99LN-BrM endpoint tumors respectively (Extended data Fig. 3d-g). We determined the time point when tumor lesions were first detectable, and evaluated effects on overall survival using the 99LN-BrM and MDA-BrM mouse models (Fig. 3j). We found a significant delay in tumor onset in mice that developed BrM in the context of BLZ945 treatment in both models (Fig. 3k,l, and Fig. 3n,o). This effect translated into significantly prolonged median survival of BLZ945-treated animals in the MDA-BrM model (Fig. 3m). However, once tumors developed, the growth rate was similar in vehicle and BLZ945-treated animals (Extended data Fig. 3h). We observed similar effects with a delay in tumor onset in the 99LN-BrM model, but no significant differences in median survival were detected (Fig. 3p, see also Extended data Fig. 3i). This indicates that tumor cells can ultimately compensate for the loss of supportive TAM functions after successful metastatic colonization either by tumor cell-intrinsic traits or by engaging the support of other immune or stromal cell types that are not CSF1R-dependent.

### CSF1R inhibition transiently blocks tumor growth in established BrM

We subsequently evaluated the effects of CSF1R inhibition in established BrM, and stratified tumor-bearing mice into vehicle and BLZ945 treatment groups, starting with a similar tumor volume, based on BLI output or volumetric MRI measurements. Mice were then treated once daily for 7 days or until symptoms developed. Measurements of tumor progression were performed on d4 and d7 after treatment initiation or once per week until the trial endpoint (Fig. 4a). Imaging by BLI or MRI revealed that TAM depletion leads to a blockade of tumor outgrowth in the majority of BLZ945-treated animals, with a few animals showing regression of large BrM lesions (Fig. 4b-e). Quantification of the depletion rate indicated 63% or 65% decrease of IBA1+ TAMs in MDA-BrM or 99LN-BrM respectively (Extended data Fig. 4a-d). Flow cytometry further confirmed depletion of CSF1R-expressing cell types including MG, MDM and inflammatory monocytes, while the relative abundance of granulocytes increased (Extended data Fig. 4e,g). Other tumor-associated cell types including T cells in the 99LN-BrM model were not affected by CSF1R inhibition (Extended data Fig. 4f,h). Decreased tumor burden was associated with a reduction in the proliferation:apoptosis (Ki67:CC3) index in BLZ945-treated MDA-BrM and 99LN-BrM lesions (Extended data Fig. 5a-d). However, while CSF1R inhibition led to initial tumor responses at d4 and d7 after treatment initiation (Fig. 4d,e, see also Extended data Fig. 5e,f), BrM regrowth was subsequently detected in both models (Extended data Fig. 4e,f). Consequently, no significant effects on median survival in either BrM model were observed (Fig. 4f,g).

**Fig. 4.**
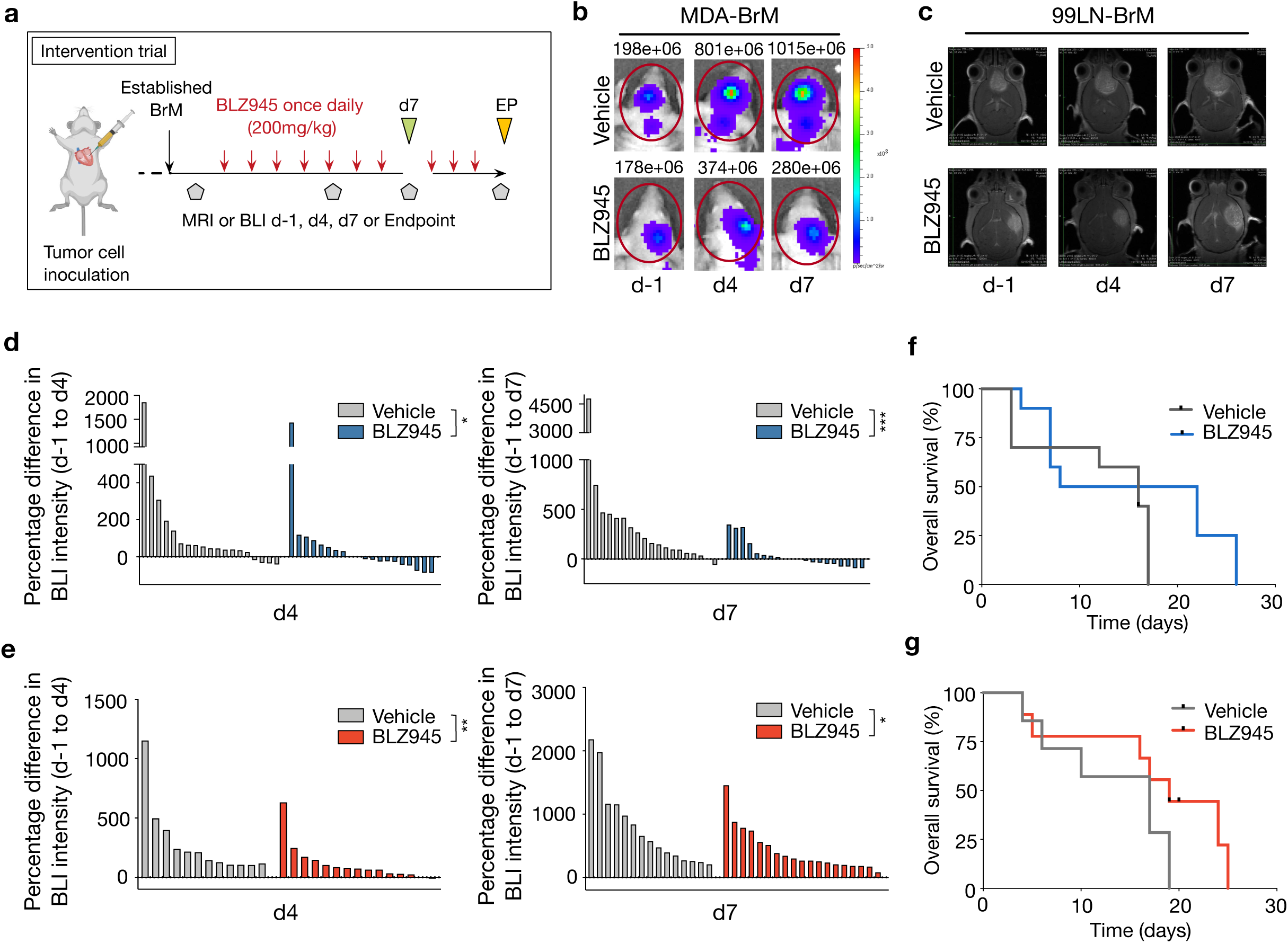
CSF1R inhibition leads to tumor stasis in short-term trials followed by rapid tumor regrowth. **a,** Experimental design of intervention trials. **b,** Representative BLI images of vehicle and BLZ945-treated MDA-BrM mice. **c,** Representative MRI images of vehicle and BLZ945-treated 99LN-BrM mice. **d,e,** Waterfall plots of relative BrM growth at indicated time points (MDA-BrM in **(d)**: n=19, vehicle and n=20, BLZ945; and 99LN-BrM in **(e)**: n=12 and 15; vehicle and n=15 and 19; BLZ945). *\*P*<0.05, *\*\*P*<0.01, *\*\*\*P*<0.001, Mann-Whitney U log rank test. **f,g,** Kaplan-Meier survival curves of MDA-BrM and 99LN-BrM mice after treatment as indicated in **(a)** (MDA-BrM in **(f)**: n=10, vehicle and BLZ945; and 99LN-BrM in **(g)**: n=7, vehicle and n=9, BLZ945, Mantel-Cox log rank test).

### TAM depletion leads to gene expression changes in tumor cells

We next FACS-purified tumor cells from vehicle and BLZ945-treated MDA-BrM mice 7 days after treatment initiation to perform RNAseq for further mechanistic insight into indirect effects on tumor cells as a consequence of CSF1R inhibition (Fig. 5a,b). Analysis of DEG in tumor cells revealed that the down-regulated genes were predominantly associated with DNA repair, cell cycle or transcriptional regulation, e.g. *XRCC2, CDK1, CCNF, TOP2A* and *ESCO2,* or represented receptor tyrosine kinases involved in wound repair, tumor growth and invasiveness, e.g. *DDR2* (Fig. 5c and Extended data Table 5). Gene ontology analysis further confirmed that gene sets related to cell cycle regulation and proliferation were found among the top 20 pathways in the gene set overrepresentation analysis (ORA, Fig. 5d) of genes downregulated in MDA-BrM upon BLZ945 treatment.

**Fig. 5.**
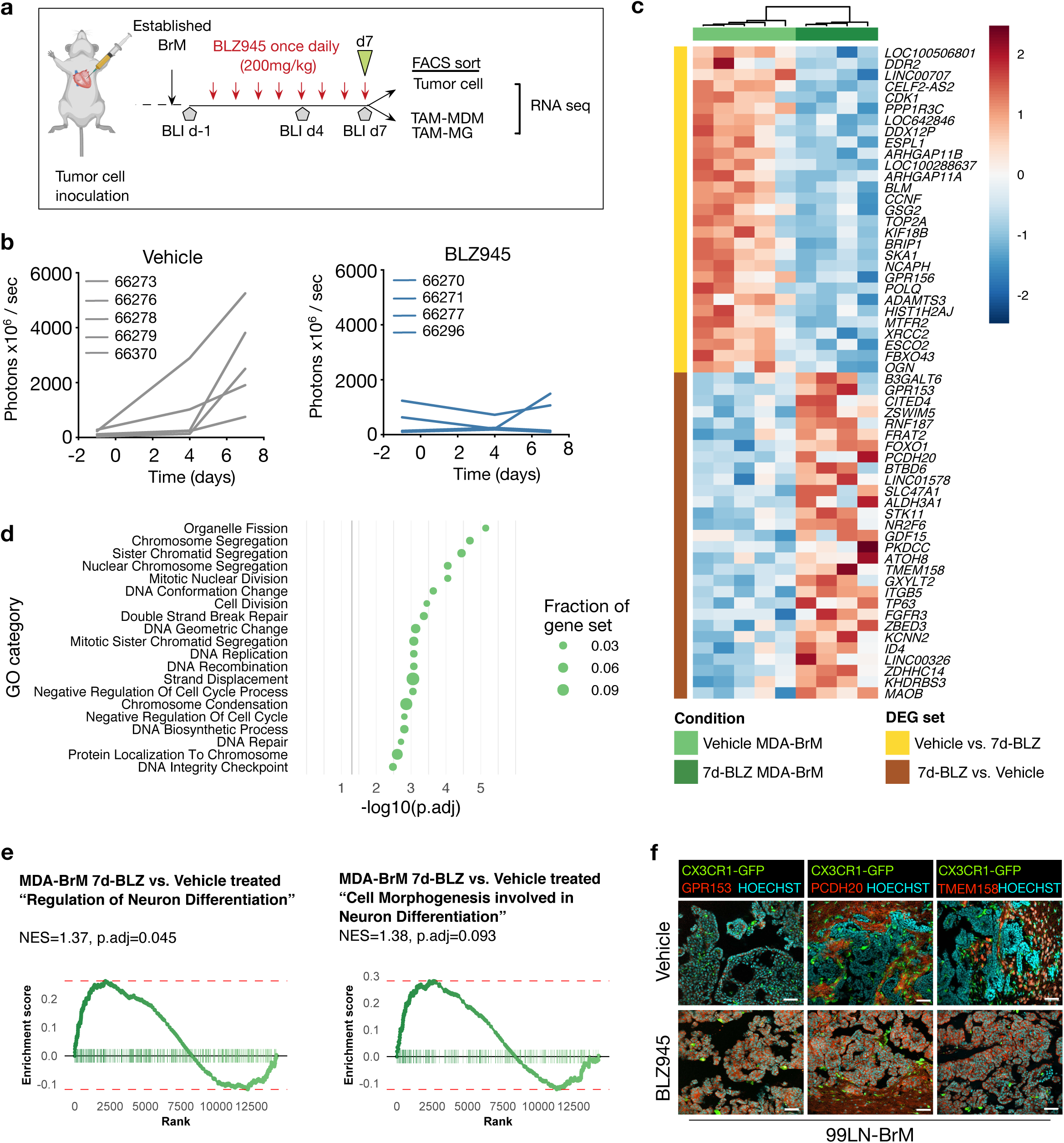
TAM depletion leads to cell cycle arrest and induction of neuronal signatures in tumor cells. **a,** Experimental design for the generation of FACS sorted samples for RNA sequencing. **b,** Quantification of BLI intensity in vehicle and BLZ945 treated MDA-BrM tumors subjected to RNAseq analysis (n=5; vehicle, n=4; BLZ945). **c,** Heatmap of DEG in MDA-BrM tumor cells in response to 7 days of BLZ945 treatment *in vivo* (n=5; vehicle, n=4; BLZ945, p-adjust≤0.1). Columns were clustered using Ward’s method with 1 – Pearsońs correlation coefficient as the distance measurement. Row annotation indicates comparison with differential expression. **d,** Top 20 (i.e. with lowest p-adjusted) pathways (from the MSigDB GO collection) identified by gene set ORA in the DEG set Vehicle treated vs. 7d-BLZ. **e,** Gene set enrichment analysis (GSEA) for signatures “Regulation of Neuron Differentiation” and “Cell Morphogenesis involved in Neuron Differentiation” in 7d BLZ945 treated tumor cells compared to vehicle controls. NES, normalized enrichment score. **f,** Representative images of tumor cell-expressed factors that are associated with neuronal differentiation identified in panel **(c)**. Scale bars, 50 μm.

Interestingly, evaluation of the list of up-regulated DEG revealed multiple genes that are typically enriched in cells of the CNS, e.g. *GPR153*, or are associated with diverse CNS pathways (Fig. 5c). Among these *PCDH20*, *BTBD6* and *TMEM158* are known to be involved in CNS cell-cell communication, neuronal differentiation, or neuronal survival, respectively ^31–34^. Gene set enrichment analysis (GSEA) additionally demonstrated expression changes in genes annotated as ‘Regulation of Neuron Differentiation’ (GO0045666) and ‘Cell Morphogenesis involved in Neuron Differentiation’ (GO0048812) in tumor cells following *in vivo* BLZ945 treatment (Fig. 5e). Histological evaluation confirmed higher expression of GPR153, PCDH20 and TMEM158 in tumor cells in BLZ945-treated animals in the 99LN-BrM model (Fig. 5f). Together these data suggest a form of neuronal mimicry, which may represent a general trait of brain metastatic tumor cells to integrate into CNS cell-cell communication circuits that support cell survival-in this case, as an adaptation to TAM depletion.

### CSF2Rb-STAT5 downstream signaling compensates for CSF1R inhibition in TAMs

To gain functional insight into the direct effects of CSF1R inhibition on TAMs, we FACS-purified TAM-MG and TAM-MDM from vehicle and BLZ945-treated animals (Extended data Fig. 6a-e, Extended data Table 3), as well as MG and blood monocytes from normal mice as controls for RNAseq analyses. We first queried the expression of putative M1-like and M2-like genes ^35^ and found that TAMs acquire a mixed phenotype in BrM based on expression of polarization markers and transcription factor activity (Fig. 6a,b). In contrast to TAMs in glioblastoma ^19,21^, we did not observe a loss of M2-polarization of TAMs in response to CSF1R inhibition. To further interrogate the consequences of CSF1R inhibition on TAM activation states, we performed gene set ORA to identify pathways that are enriched in BLZ945-treated TAM-MG and TAM-MDM compared to the vehicle-treated TAM populations, when contrasted to their normal counterparts isolated from tumor-free mice (Extended data Fig. 6f). BLZ945-treated TAMs showed an enrichment of gene sets that are associated with neuro-inflammation typically found in neurological disorders including EAE/MS or Alzheimer’s disease, such as TNF and IL6 (Extended data Fig. 6f) ^36,37^. Gene signatures associated with metabolic modification and cellular responses that indicate the production of nitric oxide (NO) and reactive oxygen species (ROS), were also elevated (Extended data Fig. 6f). Collectively, these gene ontology analyses revealed the presence of a signaling network, which includes inducer- and effector molecules of neuro-inflammation.

**Fig. 6.**
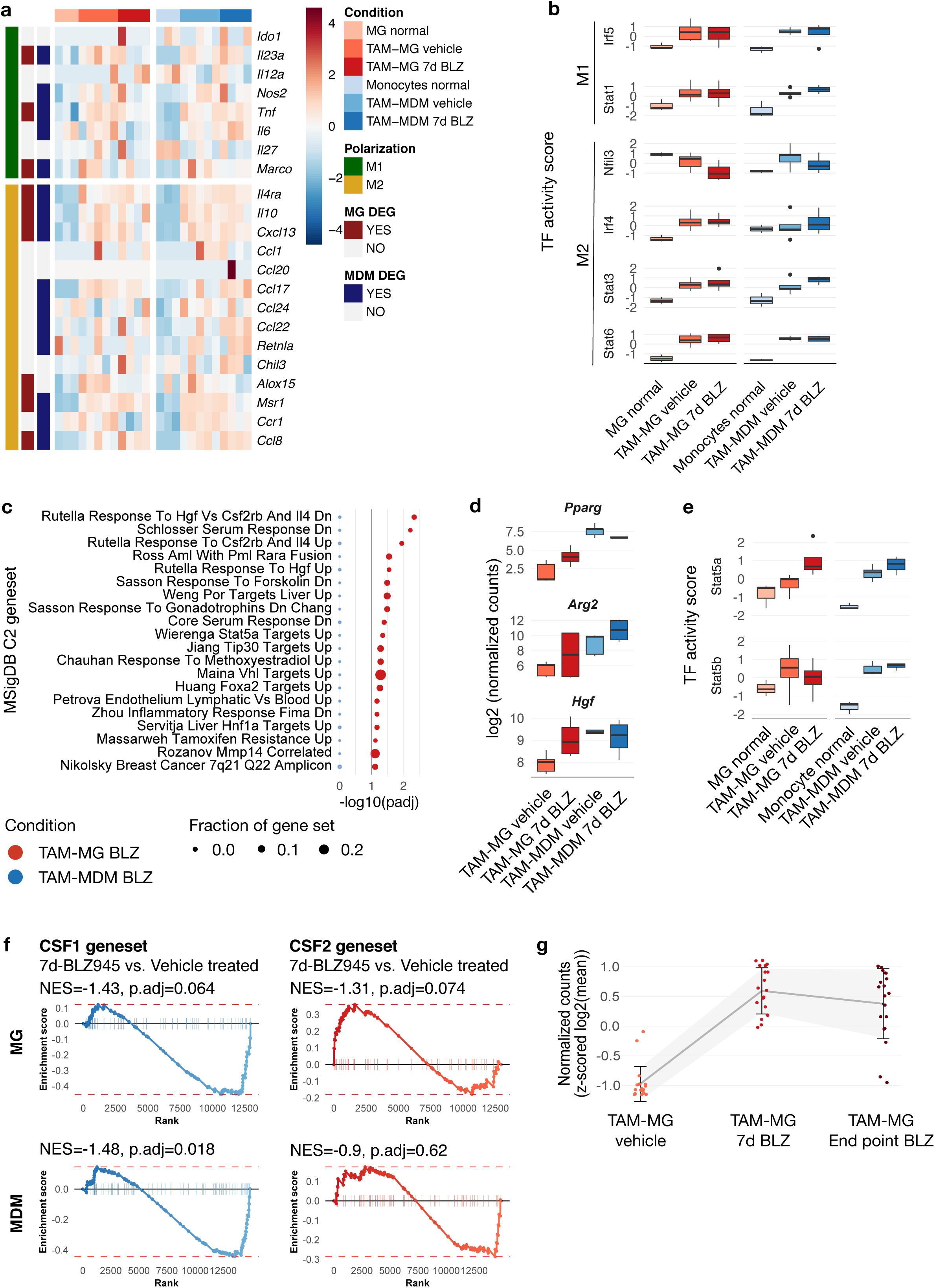
CSF1R inhibition induces compensatory CSF2-mediated activation of TAMs. **a,** Expression heatmap of M1- and M2-like macrophage markers in vehicle and 7d BLZ945 treated TAMs (n=5; vehicle, n=4; BLZ945) from the MDA-BrM model and control monocytes or normal MG (n=3 each). **b,** Transcription factor (TF) activity for a selected panel of M1/M2 associated transcription factors. **c,** Gene set ORA to identify pathways (top 20 based on their p-adjust, from the MSigDB C2 collection) that are induced in 7d-BLZ945 vs. Vehicle treated TAMs in MDA-BrM tumors (n=5; vehicle, n=4; BLZ945, fdr≤0.1). **d,** Box plots of CSF2 target gene expression and **e,** normalized TF activity scores for Stat5a and Stat5b in vehicle- and BLZ945-treated TAMs and normal MG and blood monocytes. **f,** GSEA of CSF1 and CSF2 signatures published in ^39^ analyzed in 7d BLZ945-treated TAM-MG compared to vehicle. NES, normalized enrichment score. **g,** Z-scored normalized counts of CSF2 signature genes in TAM-MG in vehicle-, 7d- and end point-BLZ945 treated MDA-BrMs.

We then investigated putative signaling pathways mediating this phenotype upon CSF1R inhibition by directly comparing the expression of the subset of TAMs that are protected from BLZ945-induced cell death (∼35% of cells, Extended data Fig. 4a,b) to their vehicle-treated counterparts (Fig. 6c). Our data indicate a shift from CSF1R to CSF2Rb-STAT5 downstream signaling, as an unbiased ORA analysis of the DEGs revealed an overrepresentation of CSF2Rb- and STAT5-associated pathways in BLZ945-treated MG (Fig. 6c). The induction of a pathway associated with production of MMP14 (Fig. 6c), a protease that has been correlated with demyelination in neuro-inflammation ^38^, was also observed. Moreover, our data further point towards the induction of IL4-driven pathways (Fig. 6c, see also Extended data Fig. 6f) that were associated with wound healing processes in recurrent glioblastoma in the context of long-term CSF1R inhibition ^20^. Activation of CSF2R downstream signaling was further confirmed by elevated expression of known CSF2R downstream genes and transcription factor activation (Fig. 6d,e).

To further validate that CSF1R blockade leads to induction of a compensatory CSF2Rb-STAT5 signaling axis, we compared the gene expression in TAMs from vehicle and BLZ945-treated animals to published signatures of CSF1 and CSF2 downstream signaling in MG ^39^. GSEA confirmed that in particular 7d-BLZ945-treated TAM-MG in BrM express signatures of CSF2 activation while vehicle-treated TAM-MG showed signatures associated with CSF1 activation (Fig. 6f). Gene signatures indicative of compensatory CSF2Rb signaling remained evident in samples isolated from mice that were treated with BLZ945 longer than the 7d period-denoted as BLZ endpoint (EP) (Fig. 6g). In contrast to TAM-MG, TAM-MDM lacked the CSF2 gene signature enrichment in response to CSF1R inhibition *in vivo* (Fig. 5f), although both cell types respond to CSF2 *in vitro* (Extended data Fig. 7a,b), express *Csf2ra* and *Csf2rb* (Extended data Fig. 7c), and show up-regulation of putative CSF2 target genes and STAT5 transcription factor activity in response to CSF1R inhibition (Fig. 6d,e). Taken together, our data indicate that CSF1R inhibition leads to the induction of a compensatory CSF2Rb-STAT5 signaling axis that protects a subset of TAMs from BLZ945-induced cell death in BrM. However, compensatory CSF2Rb-STAT5 activation induces TAM gene signatures that have been associated with neuroinflammation and which trigger IL4-mediated wound repair mechanisms (summarized in Extended data Fig. 7d).

### Tumor-associated niche cells are the primary source of CSF2 in BrM

We next sought to identify the cellular source of CSF2 in BrM. Gene expression analyses indicated that TAM-MG and TAM-MDM express no or low *Csf2* levels respectively, with no change in expression levels in response to CSF1R inhibition (Extended data Fig. 7c). FACS-sorted MDA-BrM tumor cells showed low *CSF2* mRNA expression, however, we did not detect CSF2 protein in tumor cells in MDA-BrM and 99LN-BrM lesions by IF staining (Fig. 7a). Histological assessment revealed that the dominant cellular source for CSF2 in vehicle and BLZ945-treated mice are vessel-associated cells within or in close vicinity to BrM lesions (Fig. 7a), indicating the generation of a protective environment within the vascular niche. Further characterization of the CSF2 expressing cell type revealed that PDGFRb+ pericytes represent the major cellular source for CSF2 in BrM (Fig. 7b). Critically, the TAMs that were still present after 7 days of BLZ945 treatment were localized in close proximity to vessels. By contrast, the number of TAMs localized distal to vessels was greatly reduced in response to CSF1R inhibition (Fig. 7c-d). Consequently, the presence and spatial distribution of CSF2-producing cells relative to TAM-MG and TAM-MDM may determine regional differences in the extent of TAM depletion vs. CSF2-mediated activation in BrM.

**Fig. 7.**
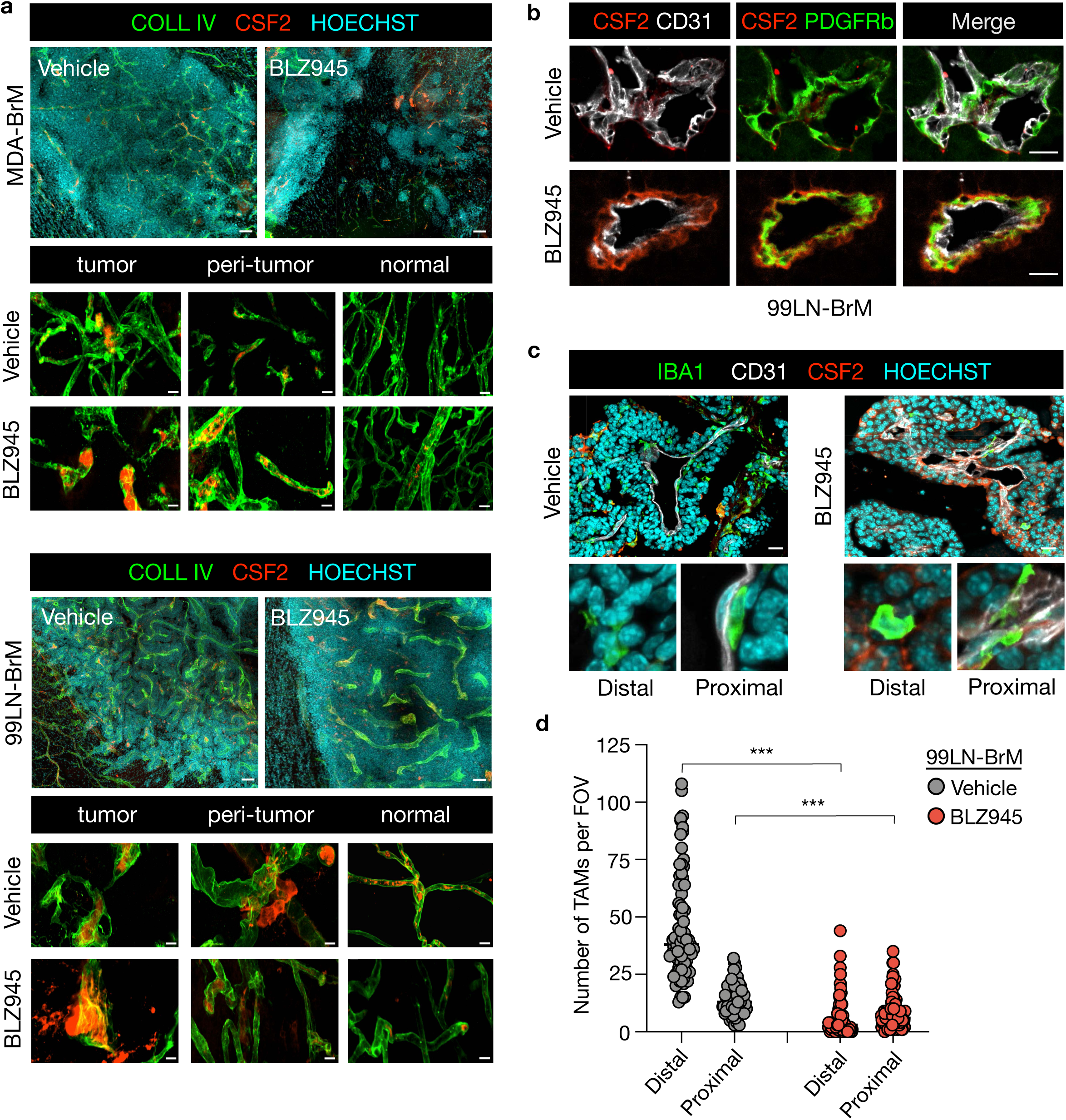
CSF2-mediated TAM activation in perivascular protective niches. **a**, Representative IF images of vehicle- and 7d BLZ945-treated MDA-BrM and 99LN-BrM brains stained for CSF2 (red) and collagen IV (COLL IV, green) to visualize vessels. HOECHST was used as nuclear counterstain. Scale bars; 50 and 10 μm. **b,** Representative IF images of vehicle- and 7d BLZ945-treated 99LN-BrM brains stained for CD31+ endothelial cells (white) and PDGFRb+ pericytes (green) in co-staining with CSF2 (red). **c,** Representative IF images of vehicle- and 7d BLZ945-treated 99LN-BrM brains stained for IBA1+ macrophages/microglia (green), CD31+ endothelial cells (white) and CSF2 (red) to visualize the localization of macrophages relative to the vascular niche. Scale bars; 25 μm. **d,** Quantification of macrophages/microglia proximal or distal to blood vessels as depicted in the representative images in vehicle and 7d BLZ945-treated animals (n=6 per group with 15-20 FOV analyzed per animal). \*\*\**P*<0.001; two-tailed Student’s t-test.

### Combined targeting of CSF1R and CSF2Rb-STAT5-mediated inflammation leads to more sustained anti-tumor responses in BrM

We next examined whether combined inhibition of CSF1R and CSF2Rb-STAT5 downstream signaling could prevent the regrowth of BrM after the initial response to BLZ945 treatment. We tested the efficacy of a neutralizing CSF2Rb antibody and the previously described STAT5 inhibitor AC4-130 ^40^ to block protective effects of CSF2 *in vitro* (Extended data Fig. 8a). We did not observe any effects on the viability of bone-marrow derived macrophages (BMDM) and EOC2 MG in response to CSF2Rb neutralization using antibody concentrations of 0.05 – 0.5 μg/ml (Extended data Fig. 8b). In contrast, we found that 5 μM and 10 μM AC4-130 blocked the proliferative and protective capacity of CSF2, while lower concentrations did not affect MG and BMDM viability *in vitro* (Extended data Fig. 8c). This was further confirmed by the dose-dependent reduction of STAT5 phosphorylation in the presence of CSF2 following treatment with 1, 5 or 10 μM AC4-130, in the presence and absence of BLZ945 (Extended data Fig. 8d). Therefore, we chose the STAT5 inhibitor AC4-130 for further *in vivo* analysis to investigate the role of compensatory CSF2Rb-STAT5 signaling in response to CSF1R inhibition.

Interestingly, the combination of AC4-130 and BLZ945 *in vivo* led to significant synergistic anti-tumor effects with reduced tumor growth kinetics until the trial endpoint (Fig. 8a-c). While we observed an initial reduction of the growth rate in AC4-130 or BLZ945 single-treated mice compared to the vehicle treatment group, monotherapy with either inhibitor was ineffective in controlling tumor outgrowth at subsequent time points (Fig. 8b,c). We observed a synergistic effect of the AC4-130 and BLZ945 combination at both d4 and d7 in MDA-BrM mice (Extended data Fig. 8e). However, symptom development resulting from extracranial lesions in this model prevented us from performing longer trials. Histological assessment revealed that BLZ945 treatment in combination with AC4-130 did not lead to more pronounced depletion of the TAM population. Instead we observed a striking change in TAM morphology (Fig. 8d, see also Extended data Fig. 8f). TAMs in the combination treatment group resembled features of ramified microglia with loss of the amoeboid morphology of activated TAMs at the tumor-stroma interface or even within the tumor area (Fig. 8d, see also Extended data Fig. 8f). Examination of MG in the adjacent brain parenchyma also indicated morphological changes. BLZ945 treatment led to retraction of protrusions and enlarged cell bodies in the remaining MG population, while combination treatment normalized this phenotype (Fig. 8d, see also Extended data Fig. 8f). AC4-130 monotherapy did not induce pronounced changes in TAM morphology or numbers.

**Fig. 8.**
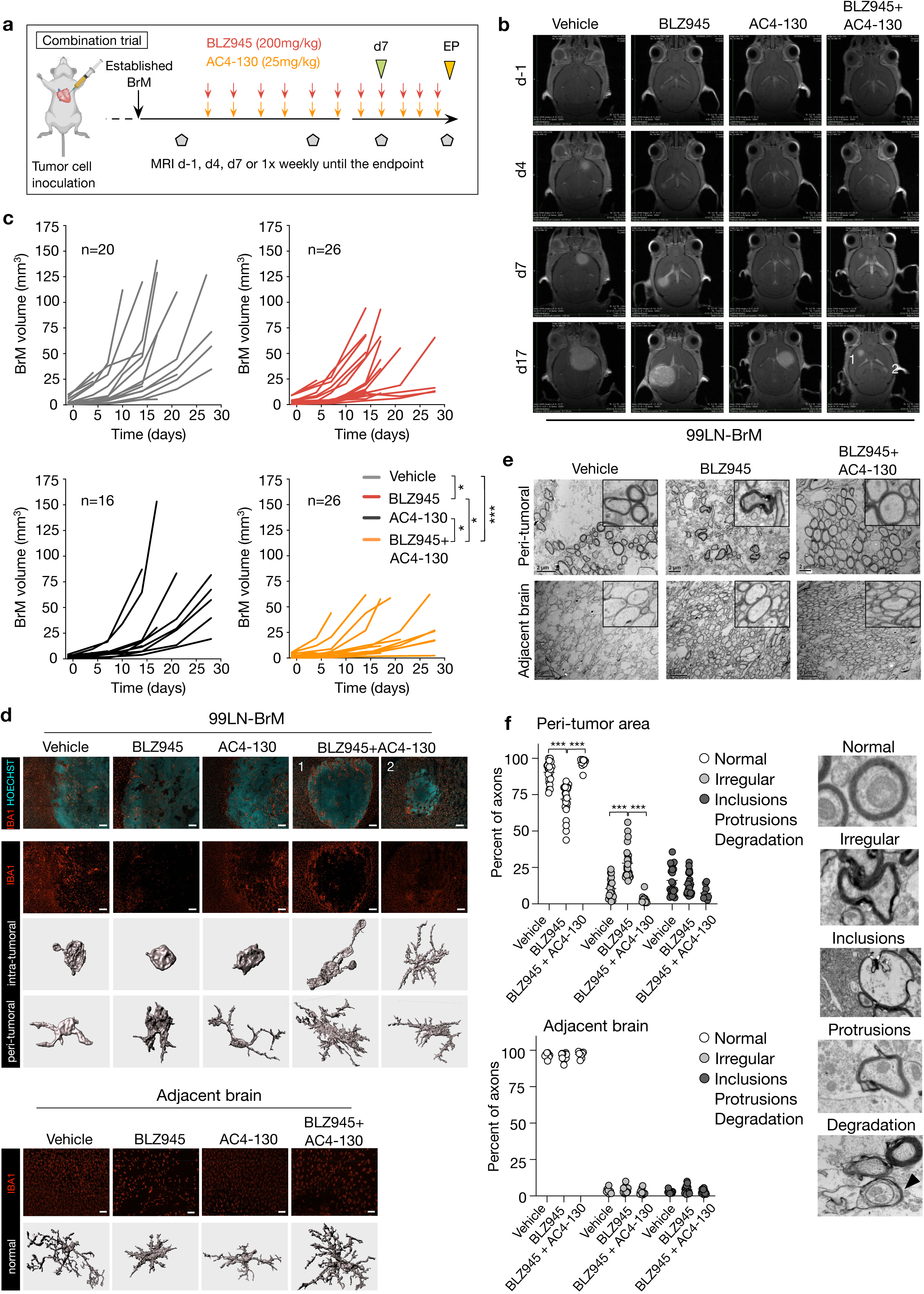
Combined CSF1R and STAT5 inhibition leads to sustained anti-tumor control and normalization of TAM morphology. **a,** Experimental design of the combination trial. **b,** Representative MRI images of 99LN-BrM mice at the indicated time points in response to treatment groups as illustrated in (a). Numbers 1,2 in the combination treatment group indicate lesions shown in panel 7d. **c,** Tumor growth curves for individual mice based on MRI volume (n=20; vehicle, n=26; BLZ945, n=16; AC4-130, n=26; BLZ945+AC4130, *\*P*<0.05, *\*\*P*<0.01, *\*\*\*P*<0.01, two-tailed Student’s t-test based on area under the curve. **d,** Representative images and 3D reconstruction of IBA1+ cells in 99LN-BrM brains. Scale bars, 100 and 25 μm. **e,** Representative transmission electron microscopy images showing axons in the peri-tumor area (upper panel) and adjacent normal brain (lower panel) in the indicated experimental groups. Scale bar, 2 μm. **f,** Quantification of the percentage of normal and abnormal axons in the indicated experimental groups in the peri-tumor area and in the adjacent normal brain (n=2 per group with 10 FOV analyzed per animal, *\*\*\*P*<0.001, two-tailed Students t-test). Abnormal axons were further classified into axons with irregular morphology (irregular) and those showing inclusions and protrusions or disaggregation of the myelin sheath indicative for degradation based on features depicted in the representative images.

We next analyzed effects on the integrity of the myelin sheath and axonal morphology by transmission electron microscopy as a mean to evaluate tissue damage following the different therapeutic regimens investigated herein, given the indications of neuro-inflammation and neuronal damage. Axons in vehicle-treated animals showed a circular morphology in the peri-tumor region similar to those found in the adjacent parenchyma. Interestingly, in BLZ945-treated BrM we observed irregularly shaped axons with dense myelin. In contrast, axons in BLZ945+AC4-130 treated BrM showed a regular morphology, indicating a reversion of the phenotype observed in response to BLZ945 monotherapy (Fig. 8e,f). We did not observe differences in other characteristics of axonal damage such as inclusions, protrusions or degradation of the myelin sheath as previously described for CSF1R haploinsufficiency ^41^. Axons in the normal adjacent brain parenchyma also showed similar morphology across all conditions (Fig. 8e,f). Collectively, our data indicate that combined targeting of CSF1R and STAT5 results in sustained tumor control, concomitant with a phenotypic normalization of the TAM and microglia population as well as reversion of the observed axonal morphology.

## DISCUSSION

Given the critical roles of various tumor-associated immune and stromal cells, TME-targeted therapies are emerging as promising intervention strategies either in combination with standard therapy or as monotherapy ^42–45^. To date, a major focus has been directed towards developing T cell-based immunotherapies such as immune checkpoint blockade or CAR T cell therapies ^46–48^. However, many cancer types are characterized by low T cell content ^44,49^, or the majority of T cells present in tumors are incapable of executing cytotoxic T cell responses ^50^. T cell-excluded tumors are often highly infiltrated by myeloid cells, which are associated with the establishment of a cancer-permissive, immune-suppressive environment ^51^. TAMs are one of the most abundant non-cancerous cell types in primary and metastatic brain tumors ^14^ and hence represent a promising target for TME-directed therapies ^16,19,20,52^. Here we demonstrate that TAM targeting with the CSF1R inhibitor BLZ945 delayed brain metastatic onset and led to temporary stasis of established metastases.

In contrast to our previous findings from glioblastoma mouse models where TAMs survived CSF1R inhibition and were instead re-educated in that context, in BrM we observed depletion of the majority of TAMs, with only a subset being protected from BLZ945-induced cell death. CSF1R inhibition triggered pro-inflammatory responses associated with the production of pro-inflammatory cytokines as well as ROS and NO. Induction of pro-inflammatory responses has been shown to be beneficial in tumor control of extracranial cancers and displays synergy with chemotherapy in a breast cancer model ^53^. By contrast, strategies that potentially initiate pro-inflammatory responses in the brain have to be very carefully weighed for their ability to achieve efficient tumor control while minimizing the risk of detrimental neurotoxicity. Indeed, our gene expression analysis revealed that the activation status of BrM-TAMs in response to CSF1R blockade resembles features of effector cells that drive demyelination and neurotoxicity in neurodegenerative disorders including EAE/MS and Alzheimer’s disease ^37^. This phenotype can be attributed to the induction of CSF2Rb-STAT5 downstream signaling in BrM-TAMs that were located in close vicinity to CSF2-producing niche cells.

Our data show that CSF1R inhibition in BrM initially induces anti-tumor responses. However, the subsequent activation of a CSF2Rb-STAT5-mediated pro-inflammatory response in the CNS most likely leads to tissue damage followed by wound repair mechanisms to limit neurotoxicity. While we did not find indications for demyelination, we did observe the formation of irregularly shaped axons with dense myelin sheaths following BLZ945 treatment. One potential explanation for this phenomenon could be enhanced re-myelination during neuronal regeneration in combination with reduced de-myelination capacity due to BLZ945-induced MG depletion, thereby leading to altered myelin organization and axonal compression. In addition to a shift from CSF1 to CSF2-mediated TAM activation, our data indicate the induction of IL4-mediated responses. This is in line with previous findings in recurrent glioblastoma where an IL4-mediated wound repair signature was associated with tumor recurrence ^20^. Importantly, neuronal mimicry by brain metastatic tumor cells as previously reported ^54–57^, and as we find in this study in response to CSF1R inhibition, might allow these cells to exploit host responses that aim to limit tissue damage and consequently support tumor outgrowth.

Pharmacological blockade of compensatory CSF2Rb-STAT5 downstream signaling further emphasized the biological consequences of causing pro-inflammatory responses within the CNS. Combined CSF1R and STAT5 inhibition resulted in synergistic anti-tumor effects compared to BLZ945 or AC4-130 monotherapy. In contrast to our initial hypothesis that combined blockade of CSF1R and CSF2Rb-STAT5 signaling would reduce neuro-inflammation due to a complete loss of the TAM population, we found a normalization of TAM morphology. This indicates that a combination treatment incorporating BLZ945 and AC4-130 has the potential to revert phenotypes of tumor-mediated TAM alterations with concomitant maintenance of anti-tumor activity. TME-targeted strategies that preserve physiologically essential cell types are of critical clinical significance, in particular in the brain which controls higher cognitive functions and all vital systems. In the future, it will be critical to determine compensatory pathways that remain active upon combined inhibition of CSF1R and STAT5 downstream signaling and to determine the respective TAM activation status in order to evaluate their role in anti-tumor responses.

Taken together, the data presented in this study reveal the risk of unleashing pro-inflammatory responses in BrM upon CSF1R inhibition, but also presents the first experimental evidence for a novel strategy to overcome this adaptive resistance mechanism. By designing rational combination therapies to disrupt tumor-glial communication, we find that this leads to sustained tumor control in conjunction with a normalization of microglia/macrophage phenotypes rather than their depletion, which has evident clinical translational implications.

## Author contributions

FK, JAJ and LS designed experiments, FK, AS, ASB, TA, MS, KN, RRM, MG, BTE, RLB, PSZ, JZ and LS performed experiments and analyzed data, FK performed computational analysis, MEH, RTD, PNH and KP provided patient samples, FK, JAJ and LS wrote the manuscript. All authors edited or commented on the manuscript. JAJ and LS conceived and supervised the project.

## Acknowledgements

We thank X. Chen, M. Quick, K. Simpson, P. Dinse, A. Möckl, P. Gebhardt and E. De Iaco for excellent technical support and members of the Joyce lab and the Georg-Speyer-Haus for insightful discussion. We are grateful to Novartis for providing BLZ945, and Dr. Marion Wiesmann, Novartis, for critical input and discussion. We thank the GSH, Goethe University and MSKCC Core Facilities for Molecular Cytology for technical assistance, in particular Marion Basoglu for electron microscopy, and the Core Facility for Microscopy at the IMB Mainz for access to the Imaris software. Research in the lab of JAJ is supported by the Breast Cancer Research Foundation, Ludwig Institute for Cancer Research, University of Lausanne, Swiss Bridge Award, Swiss Cancer League (KFS-3390-08-2016), and Cancer Research UK, as well as research fellowships from the German Research Foundation (KL2491/1-1 to FK and SE2234/1-1 to LS), Fondation Medic (to FK), and Metastasis Research Center at MSKCC (to LS). Research in the lab of LS is supported by institutional funds from the Georg-Speyer-Haus jointly funded by the German Federal Ministry of Health and the Ministry of Higher Education, Research and the Arts of the State of Hesse (HMWK), as well as grants from the LOEWE Center Frankfurt Cancer Institute (FCI), the German Cancer Consortium (DKTK, partner site Frankfurt/Mainz), the German Cancer Aid (Max-Eder Junior Group Leader Program 70111752), the German Research Foundation (SE2234/3-1), the Beug Foundation for Metastasis Research and the Dr. Bodo Sponholz Foundation.

## EXPERIMENTAL MODELS AND SUBJECT DETAILS

### Mice

All animal studies were approved by the government committee (Regierungspräsidium Darmstadt, Germany) and conducted in the Georg-Speyer-Haus (GSH) in accordance with the requirements of the German Animal Welfare Act, or approved by the Institutional Animal Care and Use Committee of Memorial Sloan Kettering Cancer Center (MSKCC). The Cx3cr1-GFP/wildtype (wt) mouse line was generated as described previously ^58^. C57BL6, Cx3cr1-GFP/wt mice or Athymic/nude mice were purchased from NCI Frederick or Charles River Laboratories or bred within the GSH or MSKCC animal facilities.

### Primary cell cultures

Bone marrow-derived macrophages (BMDM) were derived from monocytes isolated from bone marrow of 6-8 week-old mice. Bone marrow-derived cells were differentiated into macrophages by culture in Teflon bags for 7d in DMEM containing 10% fetal bovine serum with 1% L-glutamine and 1% penicillin/streptomycin with the addition of 10 ng/ml CSF1. Primary microglia were isolated from 6-8 week-old mice using the adult brain dissociation kit (Miltenyi) combined with CD11b+ magnetic bead enrichment (Miltenyi) according to the manufacturer’s instructions. Primary microglia were used for experiments directly after the isolation without further culture on plastic unless otherwise specified. Media for primary microglia contained 1% FBS, 1% L-glutamine and 1% penicillin/streptomycin and were supplemented with 20 ng/ml IL34 and 5 ng/ml TGFβ.

### Cell lines

Brain metastatic (BrM) variants of the human breast cancer cell line MDA-MB-231 (here denoted as MDA-BrM ^59^), H2030-BrM, PC9-BrM and HCC-1954-BrM were provided by Dr. Joan Massagué, MSKCC, New York, and labeled with a triple-imaging vector (TK-GFP-Luc; TGL ^60^). The murine parental 99LN cell line was derived from a metastatic lymph node of the MMTV-PyMT breast cancer model (C57BL6/J background) and twofold selected *in vivo* for brain homing capacity as previously described ^17^, resulting in the 99LN-BrM2 variant used herein (denoted as 99LN-BrM throughout this manuscript). The murine parental TS1 and TS2 cell lines were derived from primary tumors of the MMTV-PyMT breast cancer model (FVB/n background) as previously described ^61^. The tumor cell lines MCF7, SKBR3, BT474, T47D were purchased from the ATCC. MDA-BrM, 99LN-BrM, TS1 and TS2 cell lines were maintained in DMEM containing 10% fetal bovine serum with 1% L-glutamine and 1% penicillin/streptomycin. The remaining tumor cell lines were maintained in RPMI medium containing 10% fetal bovine serum with 1% L-glutamine and 1% penicillin/streptomycin. The microglia cell line EOC2 and the monocytic cell line THP1 were purchased from the ATCC. EOC2 cells were maintained in DMEM containing 10% fetal bovine serum with 2% L-glutamine and 1% penicillin/streptomycin with the addition of 20 ng/ml IL34 and 5 ng/ml TGFβ. THP1 cells were cultured in RPMI medium containing 10% fetal bovine serum with 1% L-glutamine and 1% penicillin/streptomycin. THP1 monocytes were differentiated into macrophages by 24h incubation with 150 nM phorbol 12-myristate 13-acetate (PMA, Sigma, P8139) followed by 24 h incubation in RPMI medium. Human brain microvascular endothelial cells (HBMEC) and human astrocytes (HA) were purchased from Sciencell. HBMECs were cultured on gelatin-coated cell culture dishes, and HA on poly-L-lysine cell culture dishes in endothelial cell media (ECM, Sciencell) + 10% FBS supplemented with endothelial cell growth factors (ECGF, Sciencell).

### Clinical samples

All participants included in this study provided written consent. Formalin-fixed and paraffin-embedded (FFPE) tissue from archived brain metastases collected between 1999 and 2014 was processed as tissue microarrays (TMAs) and provided by P.N.H. and K.H.P.. Specimens were obtained from the UCT tumor bank (Goethe University, Frankfurt am Main, Germany, member of the German Cancer Consortium (DKTK) and German Cancer Research Center (DKFZ), Heidelberg, Germany). Approval for this study was conferred by the ethics committee UCT Frankfurt / Goethe University Frankfurt am Main, Germany; project numbers GS 4/09; SNO_01-12 and SNO_02-2015. The TMAs included specimens from 165 patients. Histopathological scoring of CSF1R expression was obtained from n=14/28 melanoma, n=43/62 NSCLC, n=9/9 SCLC n=26/33 breast carcinoma, n=5/5 renal cell carcinoma. Additionally, the TMA contained n=11 carcinoma samples, which were not otherwise specified, and n=10 rare tumors that were not included in the analyses performed for this study. Patient tissue for whole section immunofluorescent stainings was provided by M.E.H. and R.T.D.. The collection of tumor tissue samples at the Centre Hospitalier Universitaire Vaudois (CHUV, Lausanne, Switzerland) was approved by the Commission cantonale d’éthique de la recherche sur l’être humain (CER-VD, protocol PB 2017-00240, F25 / 99). Additional information about the clinical samples can be found in Extended data Tables 1,2 and ^62^.

### Generation of experimental brain metastasis and in vivo BLI and MRI measurements

The initiation of brain metastasis from the MDA-BrM and 99LN-BrM cell lines has previously been described ^16,17^. Briefly, for brain metastasis generation in xenografted and immunocompetent mice, 1×10^4^ MDA-BrM cells or 1×10^5^ 99LN-BrM cells were inoculated into the left cardiac ventricle of 6-8 week-old female Athymic/nude mice or 10-12 week-old C57BL6J mice respectively. For a subset of experiments, Cx3cr1-GFP/wt mice were used for microglia labeling instead of C57BL6/J wt mice. Tumor progression was monitored by weekly BLI or MRI measurements in the MDA-BrM and 99LN-BrM models respectively. For BLI measurement, mice were injected with 100 μl of D-luciferin (BioCat) subcutaneously and luminescence intensity was measured using an IVIS spectrum *in vivo* imaging system (Perkin Elmer) with images taken after 1 sec and 1 min exposure times. MR imaging was performed using a 7 Tesla Small Animal MR Scanner (PharmaScan, Bruker) with a volume coil as transmitter and a head surface coil for signal reception. Mice were injected intraperitoneally (i.p.) with 150 μl Gadobutrol (Gadovist, 1 mmol ml^−1^, Bayer) before the measurement. Data acquisition was performed using the Paravision 6.0.1 software with images being acquired in coronal planes. For T2-weighted images, a localized T2-multislice Turbo rapid acquisition with relaxation enhancement (T2 TurboRARE; TE/TR = 33ms/2500ms) was used while a T1-weighted RARE sequence (T1 RARE; TE/TR = 6.5ms/1500ms) was applied for obtaining T1-weighted images. Volumetric analysis of brain metastases was performed on MR image DICOM files using a segmentation tool in the ITK-Snap software ^63^.

### BLZ945 preclinical trials

#### In vivo blood-brain barrier transmigration assay

For *in vivo* extravasation experiments, Athymic/nude or Cx3cr1-GFP/wt mice were pre-treated for 7 days with 20% Captisol (vehicle) or BLZ945 (200mg/kg). At d0, 5×10^5^ BrM cells were injected into the left ventricle of anesthetized mice. MDA-BrM cells were labeled with cell-tracker green (CMFDA) before injection. 99LN-BrM cells were transduced with blue fluorescent protein (BFP) for visualization. For the MDA-BrM model, mice were injected with Texas Red Lectin (Biozol) to visualize the vasculature. Samples were harvested 48h after tumor cell inoculation. For the 99LN-BrM model, samples were harvested after 12, 24 and 48h and visualization of the vasculature was achieved by collagen IV staining. Analysis of extravasation was performed on cleared tissue sections as described in the image analysis section.

#### Prevention trial

For prevention trials, Athymic/nude or C57BL6/J wt mice were treated for 7 days with 20% Captisol or BLZ945 (200mg/kg). At d0, 1×10^4^ MDA-BrM cells or 1×10^5^ BrM 99LN-BrM cells were injected into the left ventricle of anesthetized mice. Mice were treated once daily with vehicle or BLZ945 until mice developed symptoms from BrM. Tumor onset and progression was monitored by weekly BLI or MRI measurements.

#### Intervention trial

For intervention trials, mice were injected with MDA-BrM or 99LN-BrM cells as above. Tumor progression was monitored by weekly BLI or MRI measurements. Once mice showed signs of established BrM defined as a BLI output >1×10^6^ photons sec^−1^ or MRI volume > 1 mm^3^, mice were stratified into vehicle or BLZ945 treatment groups at d-1 and treatment was commenced at d0 with daily doses of 20% Captisol or BLZ945 (200 mg/kg). BrM burden was evaluated on d4 and d7 after treatment initiation, followed by weekly BLI or MRI measurements. For the short-term trials, mice were sacrificed on d7 after treatment initiation. For the survival trials, animals were treated until they developed symptoms from BrM or reached a maximum volume of >100mm^3^ based on MRI measurements or 5×10^9^ photons sec^−1^ based on BLI measurements.

#### Combination trials

For combination trials, mice were injected with MDA-BrM or 99LN-BrM cells as above. Tumor progression was monitored by weekly BLI and MRI measurements. Once mice showed signs of established BrM, defined as a BLI output >1×10^6^ photons sec^−1^ or MRI volume > 1 mm^3^, the animals were stratified into vehicle, BLZ945, AC4-130 ^40^, or BLZ945+AC4-130 treatment groups at d-1 and treatment was commenced at d0 with daily doses of 20% Captisol, BLZ945 (200 mg/kg/d), AC4-130 (25 mg/kg/d) or BLZ945+AC4-130. BrM burden was evaluated on d4 and d7 after treatment initiation followed by weekly MRI measurements until the trial endpoint.

### *In vitro* blood-brain barrier transmigration assays

*In vitro* blood-brain barrier (BBB) transmigration assays were performed as previously described ^16,59^. The artificial BBB was formed with HBMECs (20,000 cells/well) in co-culture with HA cells (100,000 cells/well) on transwell inserts with fluoroblok membranes. For conditions with primary microglia, or the EOC2 microglia cell line, cells were seeded onto the astrocyte cell layer within the last hour of the 5h-seeding period. After the different cell types were seeded, the artificial BBB was allowed to form and tighten for 3 days. Cell-tracker green (CMFDA)-labeled MDA-BrM and 99LN-BrM cells were allowed to transmigrate for 18h through the artificial BBB towards a FBS gradient in the presence or absence of BLZ945 (670 nM). Tumor cell migration through empty inserts (coated with gelatin and poly-L-lysine) with or without the addition of primary BMDMs or microglia was used to determine the effect of BMDMs and microglia on tumor cell migration in the absence of an artificial BBB to determine the baseline ability of those cells to alter the migration of tumor cells. The number of transmigrated tumor cells was quantified by analyzing 25 fields of view (FOV) that were acquired with a 20x objective on a Zeiss AxioImager Z1 using TissueQuest analysis software (Tissue Gnostics) or CQ1 analysis software (Yokogawa), respectively. Analysis was performed blinded to the group allocation.

### *Ex vivo* brain slice assays

Organotypic slice cultures were prepared from 6-8 week-old Cx3cr1-GFP/wt mice. Before tissue harvest, mice were treated for 7 days with BLZ945 (200 mg/kg) to deplete microglia. Vehicle control mice were treated with 20% Captisol. Fresh brain tissue was collected from PBS-perfused mice and cut in 250 μm slices using a vibratome (Leica). Brain slices were transferred in transwell inserts in brain slice media as previously described ^64^. Brain slices were incubated at 37°C and 5% CO2 for 2 hours, before 1×10^4^ tumor cells (CXMFA+ 99LN-BrM) suspended in 1 μl media were placed on top of the brain slices. Brain slices were transferred in to the CQ1 (Yokogawa) for time lapse imaging for 18h with images taken every 15 mins at 37°C and 5% CO2. The number of contacts between tumor cells and microglia were quantified and numbers of tumor cells on brain slices were counted in 25 FOV. Analysis was performed blinded to the group allocation.

### Proliferation assays

Cell growth rate was determined using the MTS cell viability assay (Promega). Briefly, cells were plated in triplicate in 96-well plates. 1×10^3^ cells for MDA-BrM and 99LN-BrM cells, 5×10^3^ cells per well for BMDM and EOC2. For all experiments, media was changed every 48h. Cells were grown in the presence of 67 and 670 nM BLZ945 or 1, 2, 5 and 10 μM AC4-130, or 0.05, 0.1, 0.25 and 0.5 μg/ml CSF2R blocking antibody (RnD Biosystems). BMDM and EOC2 were supplemented with recombinant CSF1 or IL34+TGFβ respectively, unless otherwise indicated. For testing the efficacy of CSF2Rb or STAT5 inhibition, BMDMs or EOC2 were supplemented with CSF2 unless otherwise indicated. For BMDMs and EOC2 stimulated with tumor cell-conditioned media (Tu-CM), recombinant growth factors were not added to the media. Reduction of the MTS substrate was detected by colorimetric analysis using a plate reader according to the manufactureŕs instructions. 20 μl of MTS labeling reagent was added to each well. Absorbance was measured after 2 hours at 490 nm and 650 nm on a SpectraMax 340pv plate reader (Molecular Devices).

### RNA isolation, cDNA synthesis and quantitative real-time PCR

RNA was isolated with Trizol, DNase treated, and 1 μg of RNA was used for cDNA synthesis. The following Taqman assays were used for qRT-PCR: *CSF1R* Hs06911250_m1 and *Csf1r* Mm01266652_m1. Assays were run in triplicate and expression was normalized to Ubiquitin C (*UBC* Hs00824723_m1 and *Ubc* Mm02525934_g1) for each sample.

### Protein isolation and Western blotting

For analysis of phosphorylated STAT5 in BMDMs and the MG cell line EOC2, cells were cultured in the absence of growth factors for 12h. Cells were subsequently stimulated for 5 min with DMEM medium supplemented with CSF1, IL34 or CSF2 (RnD Systems, 10 ng ml^−1^) or Tu-CM. Whole protein lysates were isolated with RIPA buffer containing 1x complete Mini protease inhibitor cocktail (Roche) and 1x PhosphoSTOP phosphatase inhibitors cocktail (Roche). STAT5 activation was compared between vehicle and BLZ945-treated (670 nM) BMDMs and EOC2 using antibodies specific to the Tyr694 of STAT5 relative to STAT5 levels. Details about antibodies used for immuno-blotting can be found in the Extended data Table 6.

### Tissue preparation and immunostaining

Tissue for frozen histology was fixed in 4% PFA overnight and subsequently transferred into 30% sucrose until the tissue was fully equilibrated. Tissues were then embedded in OCT (Tissue-Tek) and 10 μm cryostat tissue sections were used for subsequent analyses. For immunofluorescence staining, frozen sections were thawed and dried at room temperature and rehydrated. For standard staining protocols, tissue sections were blocked in 3% BSA+0.1% Triton X100 in PBS for 1h at room temperature, followed by incubation with primary antibodies in 1.5% BSA overnight at 4°C. An additional blocking step with the mouse on mouse blocking kit (MOM, Vector Laboratories) was performed for 1h at room temperature for primary antibodies derived from mouse. Primary antibody information is listed in the Key Resources Table. Fluorophore-conjugated secondary antibodies were used at a dilution of 1:500 in 1.5% BSA in PBS for 1h at room temperature. Consecutive staining protocols were performed if two primary antibodies were derived from the same species.

Paraffin-embedded patient sections were processed using a Leica Bond Max automated staining device. The automated deparaffinization / rehydration, citrate buffer-based antigen retrieval, and blocking of unspecific protein binding and endogenous peroxidase was followed by incubation with primary antibodies (Extended data Table 6), followed by HRP labeled secondary antibodies and DAB conversion.

Patient tissue for immunofluorescence was processed as described previously ^14^. Briefly, tissue was OCT embedded by submersion in 2-methyl butane cooled with dry ice, 10 μm sections were thawed, air dried and fixed with 100% methanol. Fixed sections were rehydrated, permeabilized with PBS +0.2% Triton X-100 for 3 hours and blocked with PBS +0.5% Tween 20 +1% TSA blocking reagent (Perkin Elmer). Primary antibody (Extended data Table 6) incubations where performed overnight in the same buffer, followed by secondary antibody incubations for 1 hour. Directly-conjugated primary antibodies were used after primary and secondary antibody stainings.

For reconstruction of cellular morphology, PFA-fixed brain samples were sliced in 350 μm thick sections using a Vibratome VT1200S (Leica). Brain slices were cleared using the X-Clarity tissue clearing system (Logos Biosystem). Tissue clearing was performed at 0.6A for 3h using the X-Clarity electrophoretic tissue clearing solution. After tissue clearing, unspecific protein binding was blocked with 3% BSA in PBS containing 0.1% Triton-X100 followed by incubation with primary antibodies (Extended data Table 6) for 24h at room temperature and fluorophore-conjugated secondary antibodies were used at a dilution of 1:500 in 1.5% BSA in PBS for 12h at room temperature

### Microscopy and image analysis

Tissue sections were visualized under a Carl Zeiss AxioImager Z1 microscope equipped with an ApoTome.2 and a Tissue Gnostic stage to allow for automated image acquisition or the confocal quantitative image cytometer CQ1 (Yokogawa). Quantification of IBA1+ macrophages/microglia, GFAP+ astrocytes and the analysis of proliferation and apoptosis were performed using the TissueQuest analysis software (TissueGnostic) as previously described ^19^ or the CQ1 analysis software (Yokogawa). For 3D reconstruction, images were acquired with the CQ1 confocal microscope using the 60x objective. A range of 10 μm with 100 Z-stacks was acquired. The semi-automatic surface-rendering module in the Imaris software (Bitplane) was used to create 3D volumetric surface objects. To histologically quantify the percentage of intravascular, extravasating or extravasated tumor cells or the localization of proximal and distal macrophages/microglia, brain sections were stained for collagen IV or CD31 to visualize the vasculature. Analysis was performed blinded to the group allocation. Immunfluorescently stained clinical tissue sections were imaged with an Axio Scan.Z1 slide scanner (Zeiss) equipped with a Colibri 7 LED light source (Zeiss) using a Plan-Apochromat 20x/0.8 DIC M27 cover slip-corrected objective (Zeiss). Cell type identification and CSF1R expression analysis was performed using the VIS Image Analysis software (Visiopharm) as described previously ^14^ and analyzed within the R environment (v3.5) ^65^.

### Transmission electron microscopy

Mice were anesthetized with Ketamine/Xylazine and transcardially perfused with PBS and glutaraldehyde in 4% PFA. Fixed tissue was sectioned in 1 mm thick slices and regions of interest were chosen based on MRI images for peri-tumor areas or within the M2 cortical region with projections into the corpus callosum in the contralateral hemisphere for tumor-free brain. Brain tissue was washed twice with Na-Cacodylate buffer and post-fixation was achieved with 2% OsO4 in aqua bidest for 2h followed by two washing steps in Na-Cacodylate buffer. Fixed tissue was dehydrated in an ascending alcohol series followed by incubation in propylene oxide and infiltration overnight with a mixture of propylene oxide and araldite. Before embedding, tissue was incubated 2x 2h in pure araldite. Araldite polymerization was performed at 60°C for 48h. 1 μm thin sections were used to define the region of interest under a light microscope and images were acquired on consecutive 50 nm thin sections using a CM12 transmission electron microscope (Philips) at 4000x magnification. 5% uranyl acetate and lead citrate were used as contrast enhancers. For the quantification of normal and abnormal axons, axonal phenotypes were categorized as normal, irregular morphology, axons with protrusions or inclusions as well as axons with disaggregated myelin sheaths indicative for degradation. Analysis was performed blinded to the group allocation.

### FACS analysis and cell sorting

For blood analysis, mice were bled either via retro-orbital or submandibular routes under isoflurane anesthesia. For cell sorting, mice were anesthetized with Ketamine/Xylazine, blood collected by cardiac puncture and animals were transcardially perfused with PBS. Brain metastases were macrodissected based on *ex vivo* BLI signal or MRI images and dissociated using the Brain Tumor Dissociation Kit (Miltenyi) and a single cell suspension was generated using the OctoMACS dissociator. For non-tumor bearing controls, cerebella and olfactory bulb was removed, and the remaining cortex was dissociated using the Mouse Tumor Dissociation Kit. Cell suspensions were filtered through a 40 μm mesh filter followed by red blood cell lysis. Normal brain and brain metastases samples were incubated with Myelin Removal Beads (Miltenyi). Cell suspensions were incubated for 15 min at 4°C with FC block followed by incubation with directly conjugated antibody panels for 15 min at 4°C. Cell suspensions were (PBS + 2% fetal bovine serum) and resuspended in a DAPI solution or stained with eFluor as live-dead staining. All flow cytometry analyses were performed on a BD Fortessa device and FACS sorting was performed on an Aria III. Cells were sorted directly in Trizol LS and snap frozen in liquid nitrogen. Antibody information for the flow panels can be found in the Extended data Table 5.

### RNA sequencing and gene expression analysis

RNA was isolated by chloroform extraction and isopropanol precipitation using a glycogen carrier. RNA-sequencing libraries were generated with the SMART-Seq preparation kit (CloneTech) and fragmented with the Nextera XT kit (Illumina). Paired end, 150 base pair, sequencing was performed by Genewiz (New Jersey, USA) on an Illumina HiSeq2500. Reads were adapter trimmed and quality clipped using trim_galore (v0.4.3) in paired mode (https://github.com/FelixKrueger/TrimGalore). Trimmed reads were mapped to a concatenated mouse (mm10) and human genome (hg38) using the STAR aligner (v2.5.2b) ^66^ with default parameters. Transcript abundance was quantified using STAR with a GTF file from iGenomes (Illumina). Within each sample only species-specific read counts were retained for further analysis. A count matrix was produced in R and differential gene expression was assessed with DESeq2 (v1.22) ^67^, using a log2 fold change cutoff of +/− 1 and a false discovery rate of 5% unless otherwise indicated. Gene set overrepresentation analyses were performed with sets from the Molecular Signatures Database ^68^ using the clusterProfiler R package (v3.10.1) ^69^, while gene set enrichment analyses were done with the fgsea R package (v1.8, https://github.com/ctlab/fgsea) ^70^ and single sample gene set enrichment was assessed using the gsva R package (v1.30) ^71^. Transcription factor activity was assessed as previously described ^17^. Plots were graphed using ggplot2 (v3.2) ^72^ and heatmaps drawn with the R package pheatmap (v1.0.12, https://cran.r-project.org/package=pheatmap).

### Data and code availability

All sequencing data has been deposited to the GEO under the accession number GSE133887.

### Quantification and statistical analysis

Summary data are presented as mean ± standard error of the mean (sem), floating bars with lines indicating min, max and median, or Tukey’s box plots using GraphPad Prism software v7 or “ggplot2”. Numerical data was analyzed using the statistical tests noted within the corresponding sections of the manuscript. Statistical analyses were performed with GraphPad Prism software v7 and R (version 3.5) performing tests as indicated and were considered statistically significant, with \**P*<0.05, \*\**P*<0.01 and \*\*\**P*<0.001.

---

**Extended data Table 1.** Clinical data of the IHC brain metastasis cohort (TMAs)

| Primary tumor | Sex<br>m/f | Patient age<br>(years) |
| --- | --- | --- |
| Colon cancer<br>(n=11) | 7/4 | 56-73<br>median 66 |
| Breast cancer<br>(n=33) | 0/33 | 32-76<br>median 58 |
| RCC<br>(n=18) | 17/1 | 45-76<br>median 65 |
| NSCLC<br>(n=62) | 36/26 | 13-80<br>median 61 |
| SCLC<br>(n=9) | 4/5 | 43-72<br>median 57 |
| Melanoma<br>n=28 | 11/17 | NA |

**Extended data Table 2.** Clinical data of the IF brain metastasis cohort (whole tissue sections)

| Primary tumor | Sex<br>m/f | Patient age<br>(years) |
| --- | --- | --- |
| Breast cancer<br>(n=9) | 0/9 | 40-69<br>median 54.5 |
| NSCLC<br>(n=7) | 3/4 | 54-70<br>median 61 |

**Extended data Table 3 DEG_TAM_MDABrM**; Excel sheet Expression fold-changes in MG and MDM.

**Extended data Table 4 GO_Overrepresentation_TAM-DEG**; Excel sheet GO analysis results of clusters from Figure 2c.

**Extended data Table 5 DEG_Tumor_MDABrM**; Excel sheet Expression fold-changes in MDA-BrM vehicle-treated vs. 7d BLZ.

**Extended data Table 6** Antibodies used for histology, western blotting and flow cytometry

**Extended data Movie 1** BrainSlice_MG_vehicle; m4v file

**Extended data Movie 2** BrainSlice_MG_BLZ945; m4v file

**Extended data Fig. 1.**
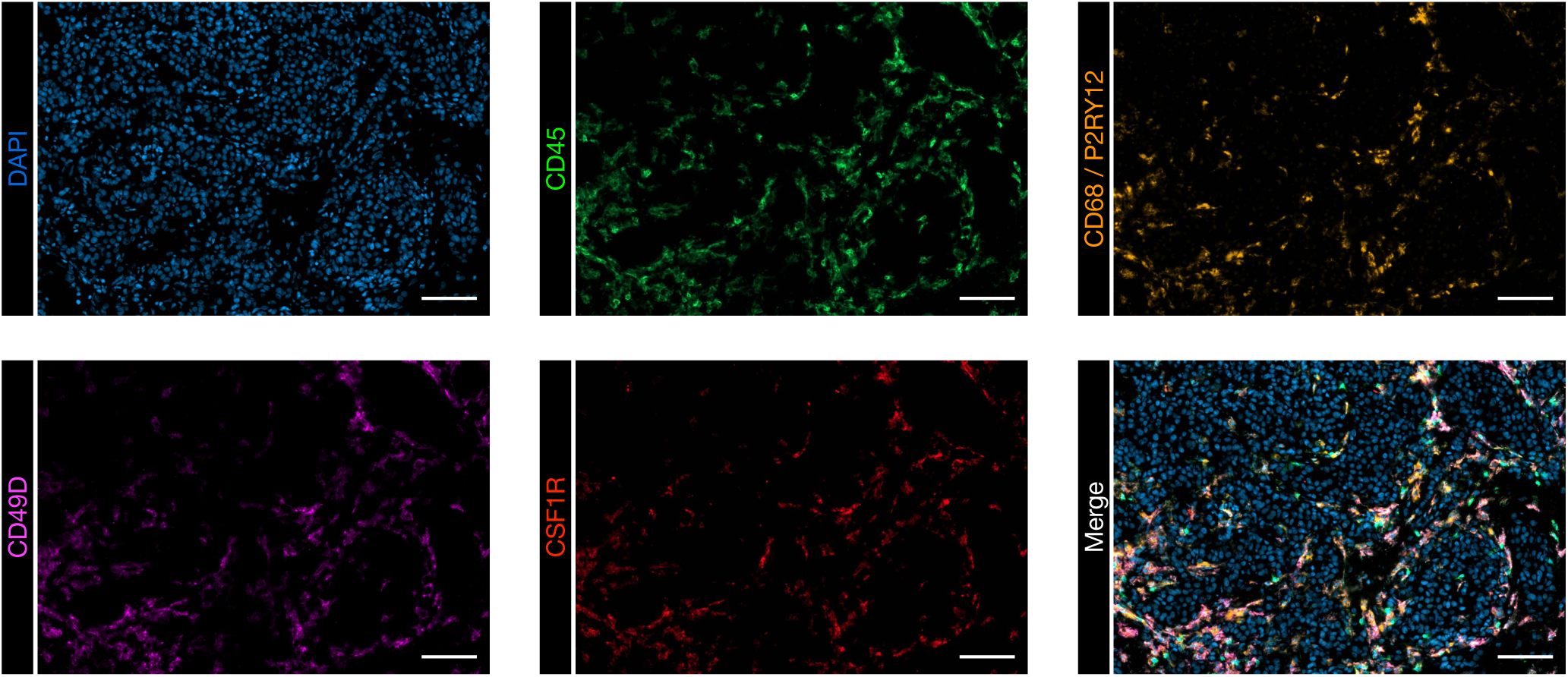
CSF1R expression in human BrM. Representative images of single channel and merged immunofluorescence (IF) images of CD45, CD68/ P2RY12, CD49D and CSF1R stainings used to identify CSF1R expression in different cell populations. Scale bar, 100 μm.

**Extended data Fig. 2.**
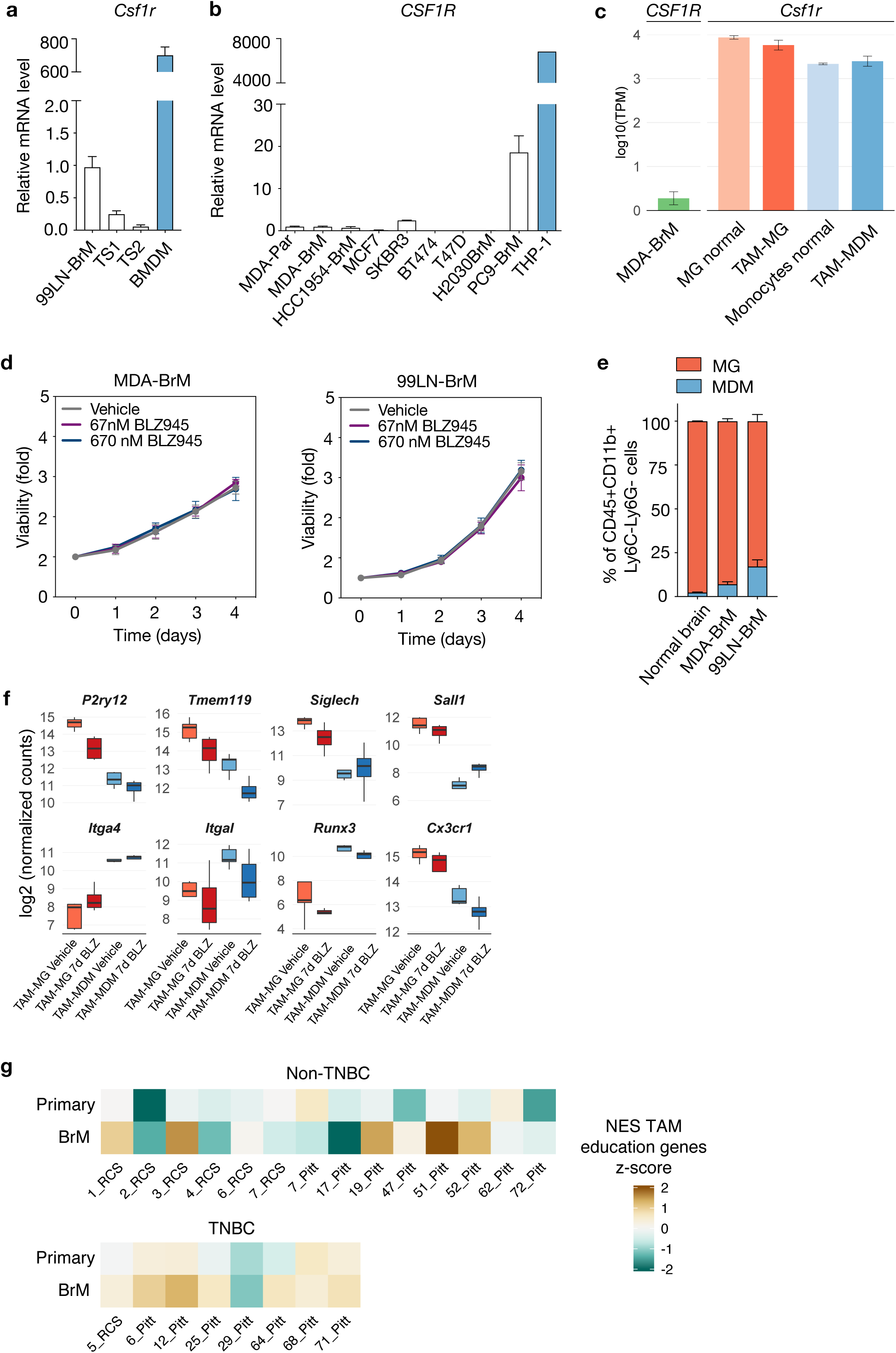
*CSF1R/Csf1r* expression in mouse BrM. **a,b,** Quantification of *CSF1R/Csf1r* expression by qRT-PCR in **(a)** mouse and **(b)** human tumor cell lines in comparison to primary bone marrow-derived macrophages (BMDM) and the macrophage cell line THP1 (n=3-9 replicates). **c,** Quantification of *CSF1R/Csf1r* expression levels by RNAseq in FACS-purified MDA-BrM tumor cells, tumor-associated microglia and macrophages (TAM-MG and TAM-MDM) as well as normal microglia and blood monocytes isolated from tumor-free mice (n=5 for tumor-bearing mice and n=3 for tumor-free controls). TPM, transcripts per million. **d,** Measurement of cell viability in MDA-BrM and 99LN-BrM cells in response to BLZ945 treatment (0 nM-vehicle, 67 nM and 670 nM) using MTS assay (n=3 replicates). **e,** Quantification of the percentage of CD49d-MG and CD49d+ MDM in the CD45+CD11b+Ly6C-Ly6G-live cell population (n=3 for tumor-free controls n=5 for MDA-BrM and n=4 for 99LN-BrM mice). **f,** Marker gene expression in normal MG and blood monocytes as well as TAM-MG and TAM-MDM to validate the purity of the FACS sorted populations. (n=5; vehicle, n=4; BLZ945). **g,** Heatmap depicting the z-standardized normalized enrichment score (NES z-score) of the union of DEG identified as BrM-TAM genes in matched human primary breast cancer and BrM from non-triple negative (Non-TNBC; upper panel) and triple negative breast cancer (TNBC; bottom panel). RNAseq data for patient breast cancer and BrM samples was obtained from ^25^. NES between primary tumors and BrM was assessed using a Wilcoxon signed-rank test (*P* = 0.021).

**Extended data Fig. 3.**
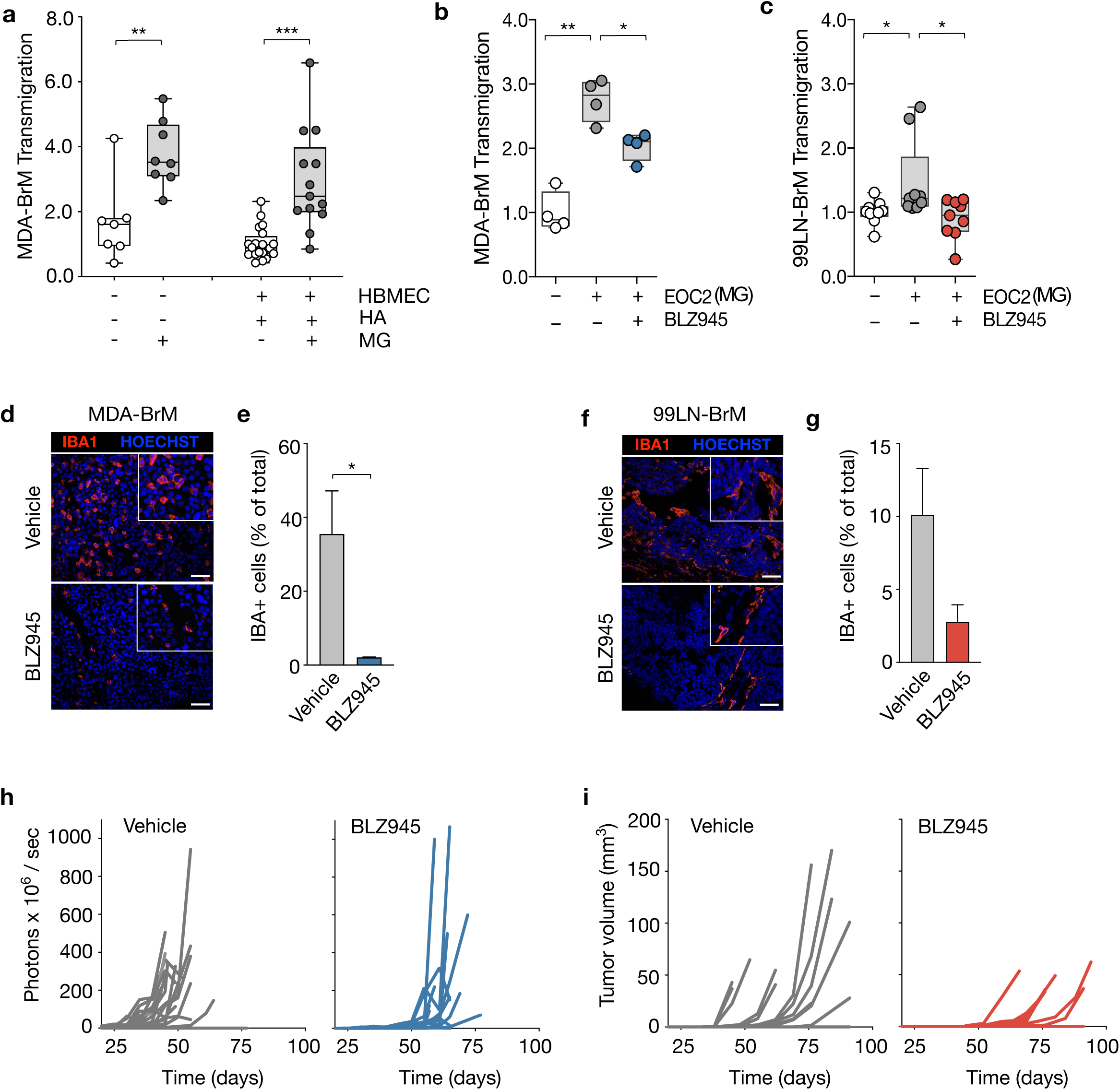
MG and BMDM support tumor cell extravasation. **a,** Quantification of the transmigration of MDA-BrM cells in the presence or absence of primary MG through an artificial blood brain barrier (BBB) formed by human astrocytes (HA) and human brain microvascular endothelial cells (HBMECs), or without an artificial BBB (n=7-11 replicates per condition). Values are relative to the control condition. **b,** Quantification of the transmigration of MDA-BrM cells, with or without the addition of the microglia cell line EOC2, in response to BLZ945 or vehicle treatment (n=4 replicates per condition). Values are relative to the control condition. **c,** Quantification of the transmigration of 99LN-BrM cells, with or without the addition of the microglia cell line EOC2, in response to BLZ945 or vehicle treatment (n=10 replicates per condition). Values are relative to the control condition. **d,** Representative immunofluorescence images of endpoint tumors from vehicle and BLZ945 treated MDA-BrM mice stained with IBA1 to visualize macrophages/microglia. HOECHST was used for nuclear counterstain. Scale bar, 50 μm. **e,** Quantification of the number of IBA1+ TAMs in vehicle and BLZ945 treated animals in MDA-BrM (n=5 for each condition). **f,** Representative immunofluorescence images of endpoint tumors from vehicle and BLZ945 treated 99LN-BrM mice stained with IBA1 to visualize macrophages/microglia. HOECHST was used for nuclear counterstain. Scale bar, 50 μm. **g,** Quantification of the number of IBA1+ TAMs in vehicle and BLZ945 treated animals in 99LN-BrM mice (n=3 for each condition). **h,** Quantification of BLI intensity during tumor progression in the MDA-BrM model in vehicle or BLZ945 treated animals in prevention trial setting (n=17; Vehicle, n=14; BLZ945). **i,** Quantification of the tumor volume based on T1 weighted MRI images during tumor progression in the 99LN-BrM model in vehicle or BLZ945 treated animals in the prevention trial setting (n=10; vehicle, n=11; BLZ945). \**P*<0.05, \*\**P*<0.01, \*\*\**P*<0.001; two-tailed Student’s t-test.

**Extended data Fig. 4.**
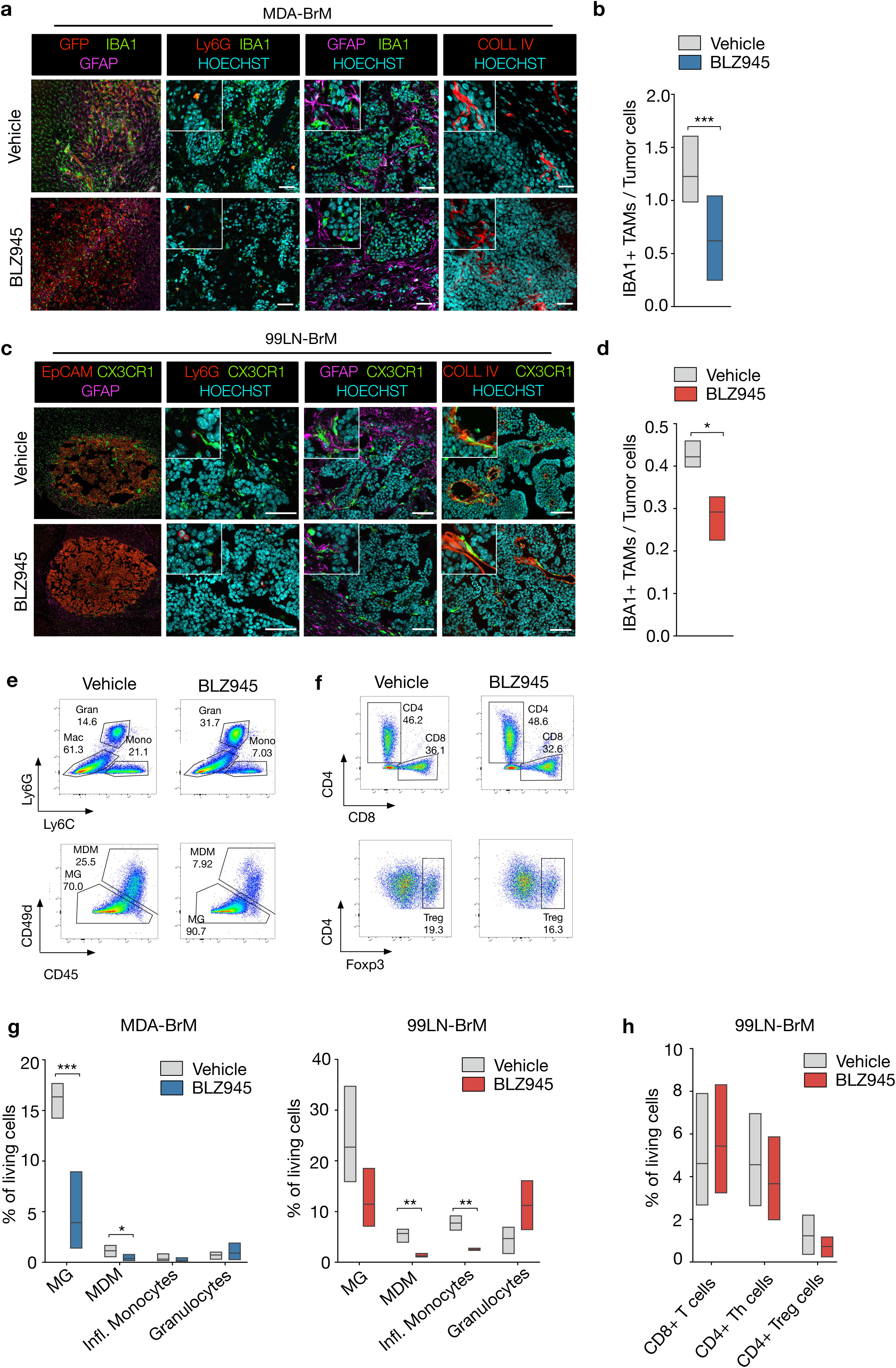
CSF1R inhibition leads to TAM depletion. **a,** Representative images of indicated markers in MDA-BrM brains after 7 days of vehicle or BLZ945 treatment. Scale bar, 50 μm. **b,** Quantification of TAMs in vehicle and BLZ945 treated MDA-BrM tumors (n=7; vehicle, n=7; BLZ945). **c,** Representative images of indicated markers in 99LN-BrM brains after 7 days of vehicle or BLZ945 treatment. Scale bar, 50 μm. **d**, Quantification of TAMs in vehicle and BLZ945 treated 99LN-BrM mice (n=3; vehicle, n=4; BLZ945). **e,** Representative FACS plots of the myeloid panel used for flow cytometric analysis. **f,** Representative FACS plots of the lymphoid panel used for flow cytometric analysis. **g,** Quantification of myeloid cell populations in the MDA-BrM (n=5; vehicle, n=4; BLZ945) and 99LN-BrM model (n=3; vehicle, n=3; BLZ945). **h,** Quantification of lymphoid cell populations in the 99LN-BrM model (n=3; vehicle, n=3 BLZ945). *\*P*<0.05, *\*\*P*<0.01, *\*\*\*P*<0.001; two-tailed Student’s t-test.

**Extended data Fig. 5.**
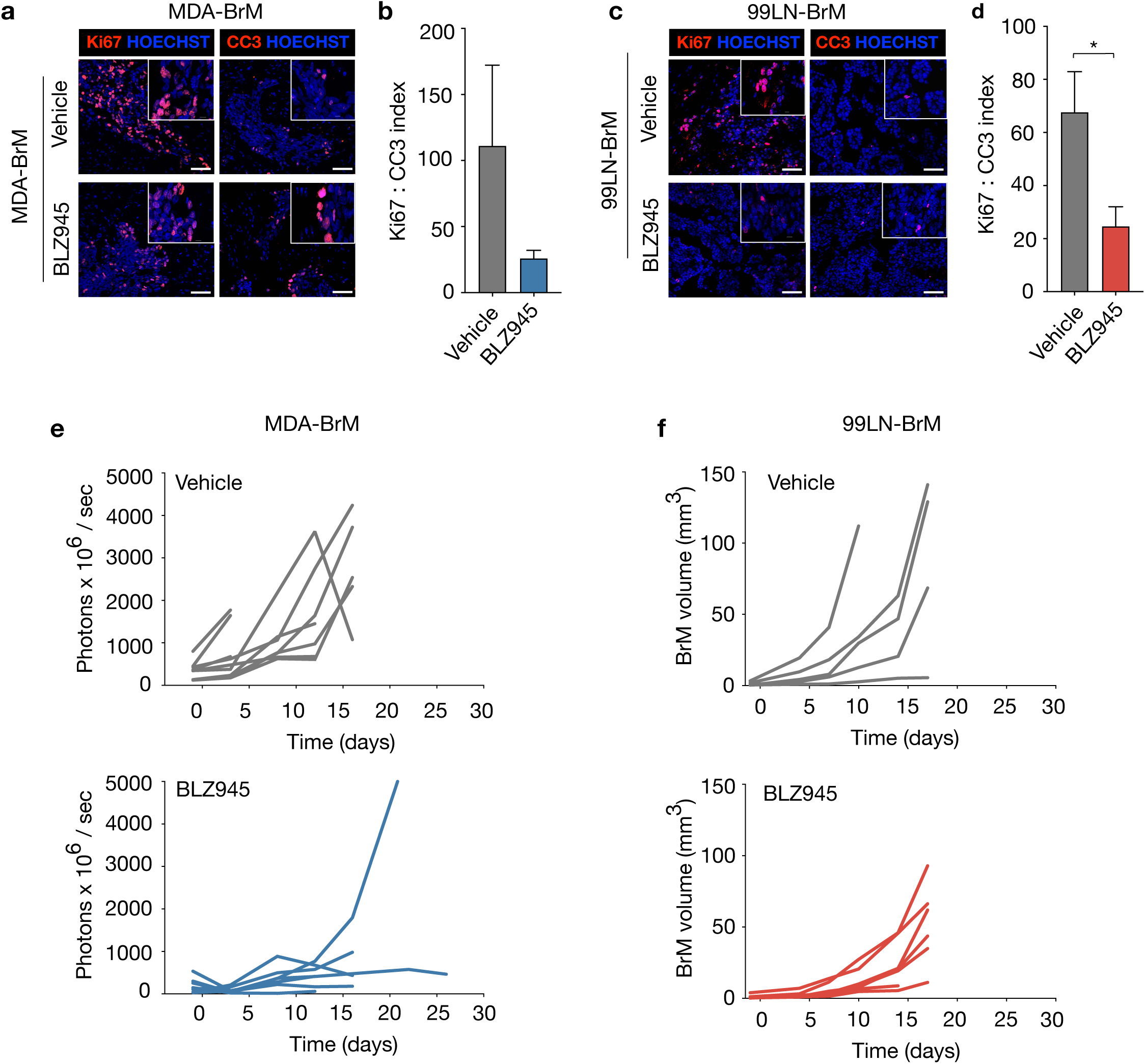
BLZ945 treatment blocks tumor growth and reduces the proliferative index of tumor cells in intervention trials. **a,** Representative immunofluorescence images stained for Ki67 (left panel) and cleaved caspase 3 (CC3, right panel) to visualize proliferating and apoptotic cells respectively in vehicle and BLZ945-treated mice from the MDA-BrM model. Scale bars, 50 μm. **b,** Ki67:CC3 proliferation:apoptosis index from immunofluorescence staining of vehicle and BLZ945-treated MDA-BrM tumors (n=5). **c,** Representative immunofluorescence images stained for Ki67 (left panel) and cleaved caspase 3 (CC3, right panel) to visualize proliferating and apoptotic cells in vehicle and BLZ945-treated mice from the 99LN-BrM. Scale bar, 50 μm. **d,** Ki67:CC3 proliferation:apoptosis index from immunofluorescence staining of vehicle and BLZ945-treated 99LN-BrM tumors (n=3). **e,** Quantification of BLI intensity of vehicle and BLZ945–treated mice from the MDA-BrM model (n=10; vehicle and n=8; BLZ945). **f,** Quantification of the tumor volume based on T1 weighted MRI images during tumor progression in the 99LN-BrM model in vehicle or BLZ945 treated animals in the intervention trial setting (n=5; vehicle, n=7; BLZ945). *\*P*<0.05; two-tailed Student’s t-test.

**Extended data Fig. 6.**
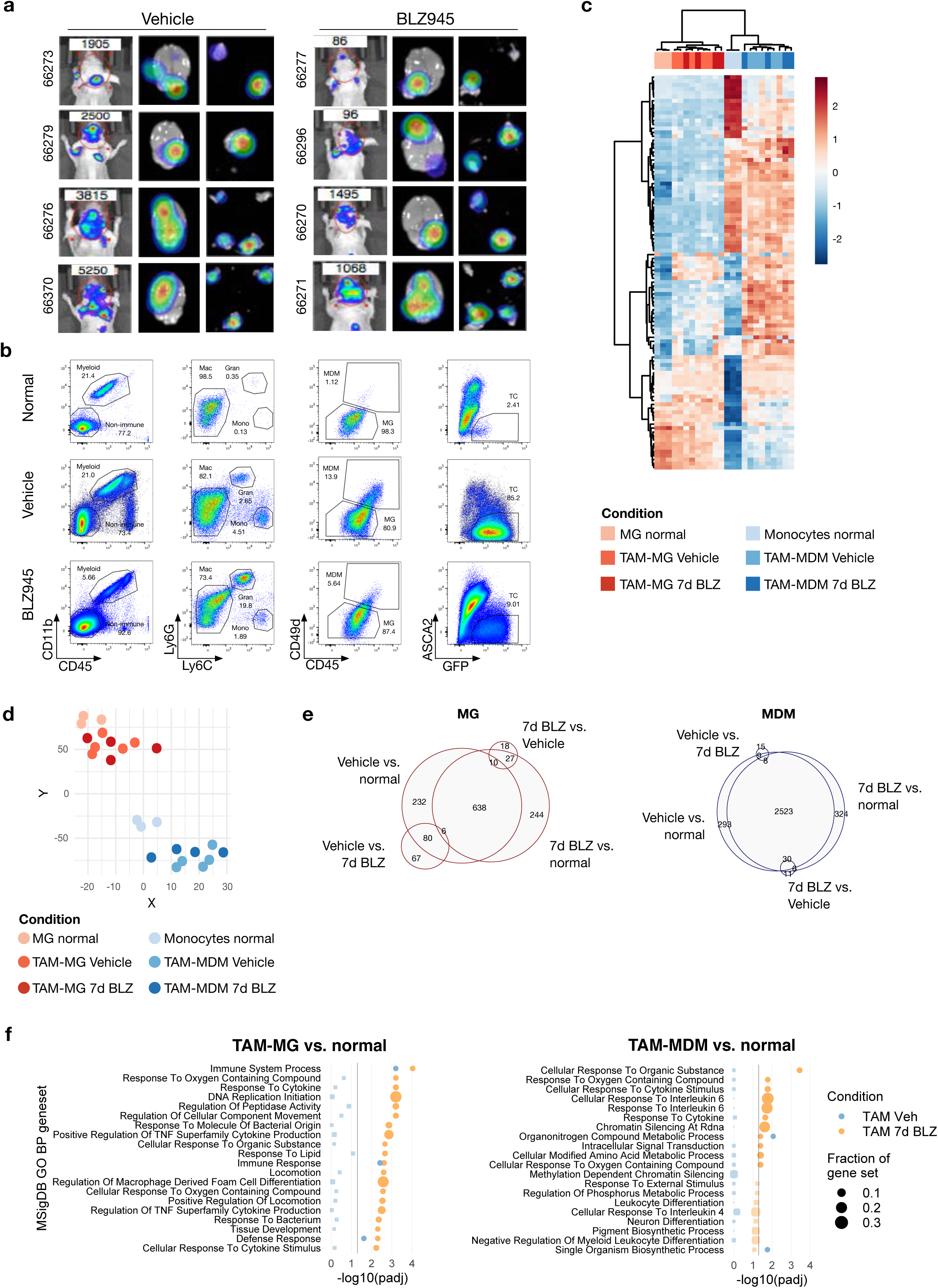
Differential gene expression in FACS-purified tumor cells and myeloid cell populations. **a,** Representative *in vivo* and *ex vivo* images of vehicle and BLZ945-treated MDA-BrM mice at d7 after treatment start. **b,** Representative FACS plots of the gating strategy for FACS purification of GFP+ tumor cells and TAM-MG (CD45+CD11b+Ly6C-Ly6G-CD49d-) and TAM-MDM (CD45+CD11b+Ly6C-Ly6G-CD49d+) from the MDA-BrM model used for RNAseq experiments. Tumor-free mice were used to isolate blood monocytes and normal MG. **c,** Heatmap of 100 most variant genes in myeloid populations across all samples. Columns and rows were clustered using Ward’s method with 1 – Pearsońs correlation coefficient as the distance measurement. **d,** tSNE plot on 2000 most variant genes shows clustering of individual samples. **e,** Euler plot depicts shared and unique DEG (p-adjusted<0.05, log2fc>1, mean expression > 10) in MG (right panel) and MDM (left panel) in the indicated comparisons. **f,** Gene set ORA (from the MSigDB GO collection) of DEGs that are specifically induced in MDA-BrM TAM-MG and TAM-MDM in response to CSF1R blockade compared to normal MG and blood monocytes (fdr≤0.1).

**Extended data Fig. 7.**
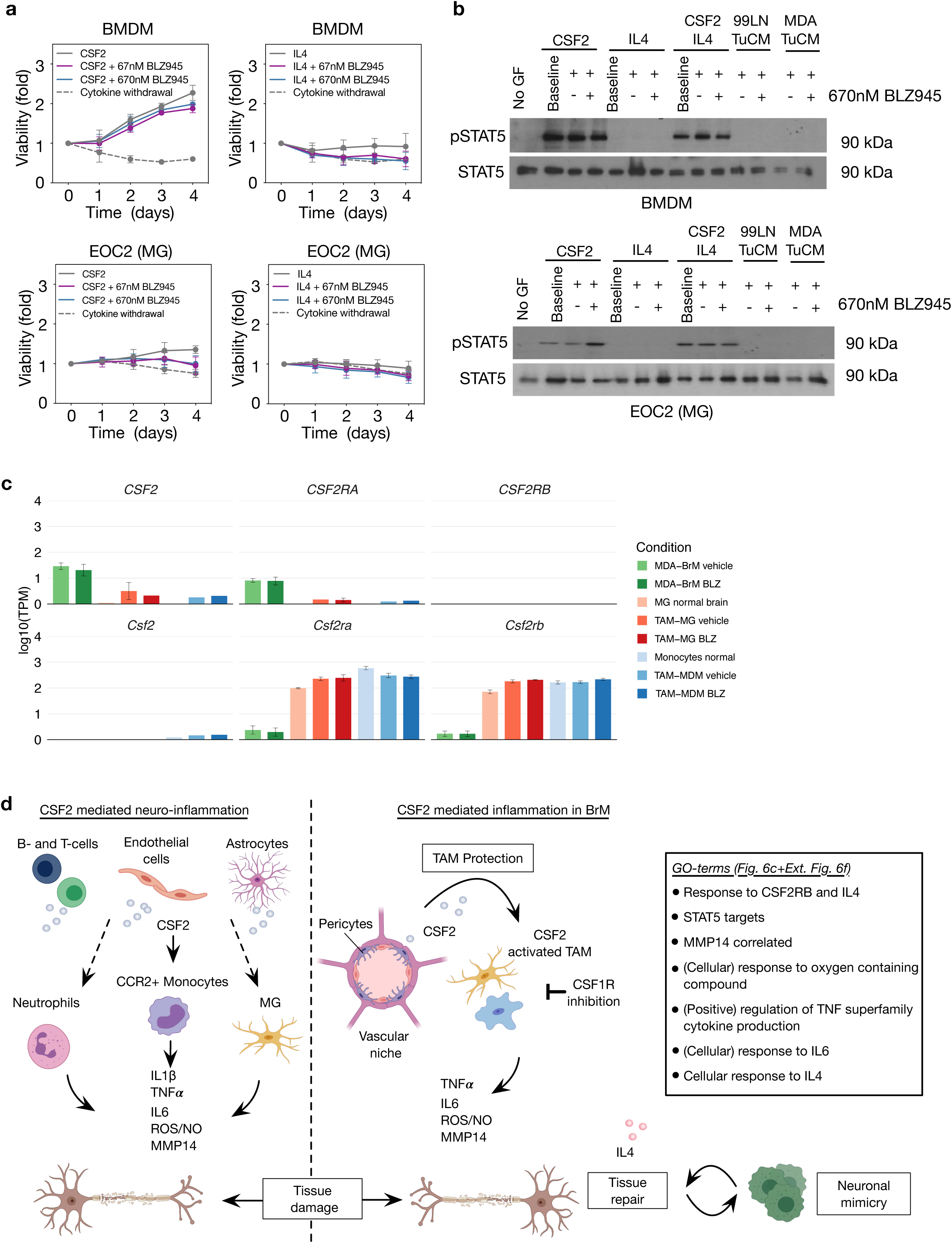
CSF2 protects TAMs from BLZ945 induced killing. **a,** Quantification of the cell viability of BMDM and EOC2 MG cells in response to CSF2 and/or IL4 stimulation in the presence or absence of 67nM and 670nM BLZ945 (n=4-8 replicates per condition). **b,** Western blot analysis of the CSF-2R downstream signaling effector STAT5 in response to cytokines or tumor cell-conditioned media (Tu-CM) stimulation in the presence or absence of 670nM BLZ945 in BMDMs and EOC2. Result shown is representative of three independent experiments. **c,** Gene expression of *CSF2/Csf2*, *CSF2RA/Csf2ra* and *CSF2RB/Csf2rb* in FACS purified MDA-BrM tumor cells, TAM-MG and TAM-MDM as well as normal MG and blood monocytes and MG (n=5; tumor-bearing mice and n=3; tumor-free controls). **d,** Model summarizing the findings from this study in comparison to results from models of neuro-inflammation. In neuro-inflammation, B- and T-cells as well as endothelial cells and astrocytes are known sources for CSF2. CSF2-activated CCR2+ monocytes are the predominant cell type in EAE models to secrete effector molecules that lead to neurotoxic tissue damage. CSF2-activated TAMs in BrM displayed gene signatures that indicate the activation of similar effector molecules (TNF, IL6, ROS/NO and MMP14) based on the listed GO terms we uncovered herein.

**Extended data Fig. 8.**
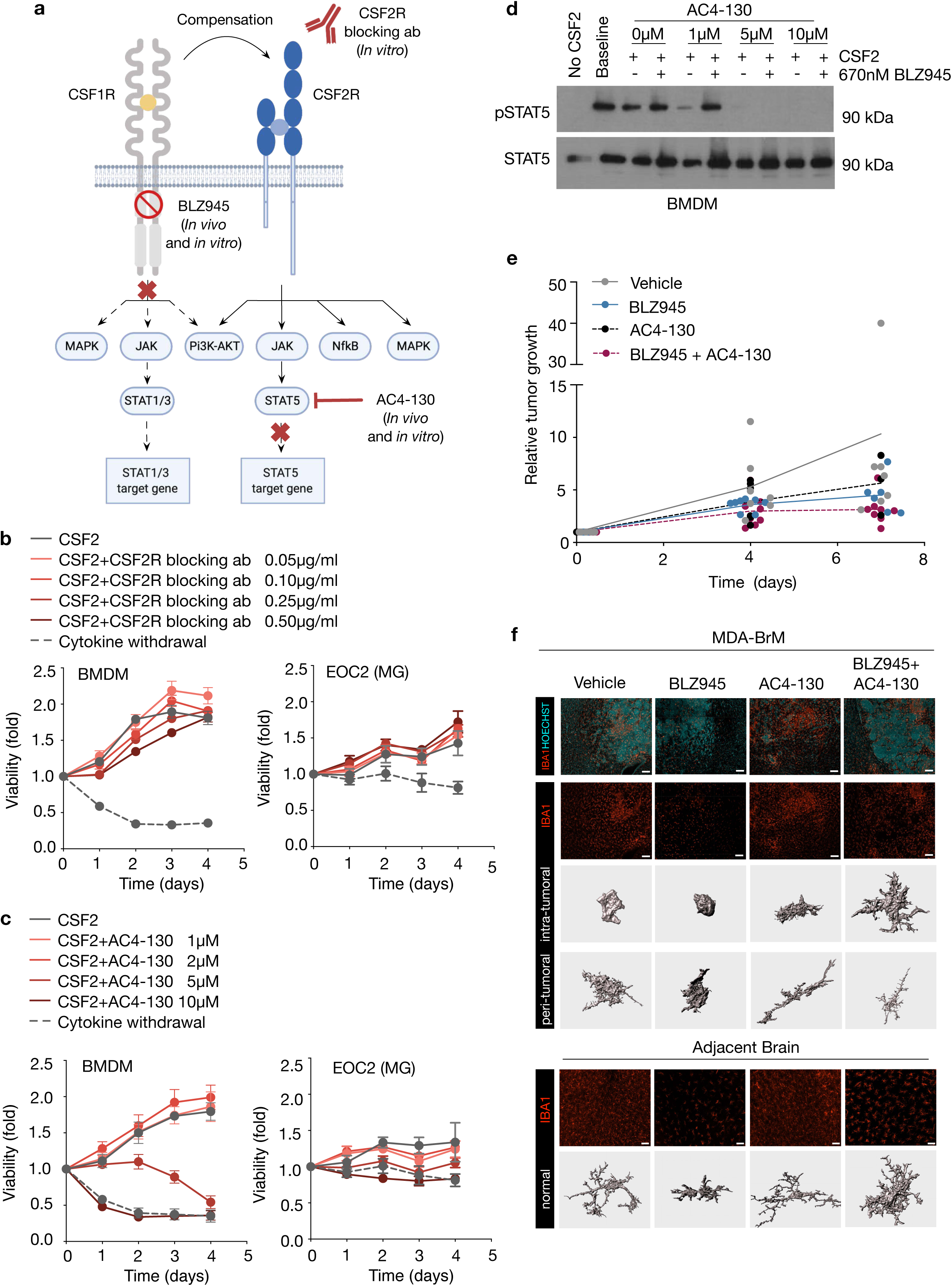
CSF1R and STAT5 inhibition *in vitro* and *in vivo*. **a,** Schematic overview of CSF1R and CSF2Rb downstream signaling. Molecular targets inhibited in this study (in preclinical models and in cell culture) are indicated. **b,** Quantification of cell viability in BMDM and EOC2 MG cells in response to increasing concentrations of neutralizing CSF2Rb antibody by MTS assay (n=4 replicates per condition). **c,** Quantification of cell viability in BMDM and EOC2 cells in response to increasing concentrations of the STAT5 inhibitor AC4-130 by MTS assay (n=4-12 replicates per condition). **d,** Western blot analysis of the CSF2R downstream signaling pathway STAT5 in response to CSF2 and increasing concentrations of AC4-130 in the presence or absence of 670nM BLZ945. Result shown is representative of three independent experiments. **e,** Quantification of the relative tumor growth rate based on volumetric evaluation of MRI images in the MDA-BrM model (n=8; vehicle, n=10; BLZ945, n=6; AC4-130, n=8; BLZ945+AC4130). Data is represented as individual replicates with lines depicting the mean of each experimental group. Analyzed by two-tailed Students t-test based on area under the curve. **f,** Representative images and 3D reconstruction of IBA1+ cells in the MDA-BrM model depicting the morphology of TAMs in intra- and peri-tumoral areas as well as normal adjacent parenchyma. Scale bars, 100 μm and 25 μm.

