## Extended data Table 6 for "Compensatory CSF2-driven macrophage activation promotes adaptive resistance to CSF1R inhibition in breast-to-brain metastasis"

| **Antibodies used for histology**  Human samples | | |
| --- | --- | --- |
| *Antigen* | *Vendor* | *Catalogue number* |
| CD45 goat anti-human | LSBio | LS-B14248-300 |
| CD49D clone PS/2 rat anti human | abcam | ab25247 |
| CD68 clone KP1 mouse anti-human | abcam | ab955 |
| CSF1R clone 9-4D2-1E4 rat anti-human AF647 | Biolegend | 347315 |
| CSF1R rabbit anti-human | Invitrogen | PA5-14569 |
| IBA1 Rabbit anti-human | Novus Biologicals | NBP22-19019 |
| P2RY12 rabbit anti-human | Sigma-Aldrich | HPA014518 |
| **Antibodies used for histology**  Mouse samples | | |
| CD31 goat anti-mouse | RnD Systems | AF3628 |
| CC3 clone 5A1E rabbit anti-mouse | Cell Signaling | 9664 |
| Csf2 mouse anti-mouse | abcam | ab54429 |
| Collagen IV rabbit anti-mouse | Merck | AB756P |
| EpCAM rabbit anti-mouse | abcam | ab71916 |
| GFAP rabbit anti-mouse | abcam | ab53554 |
| GFP chicken anti-jelly fish | abcam | ab13970 |
| GPR153 rabbit anti-human/mouse | Sigma Aldrich | SAB450166 |
| Iba1 Rabbit anti mouse | Wako Chemicals | 019-19741 |
| Ki67 clone SP6 rabbit anti-mouse | abcam | ab16667 |
| Ly6G clone RB6 8C5 rabbit anti-mouse | abcam | ab25377 |
| NeuN clone ERP12763 rabbit anti-mouse | abcam | ab177487 |
| PCDH20 rabbit anti-human/mouse | Novus Biologicals | NBP2-30010 |
| PDGFR beta rabbit anti-human/mouse | abcam | ab32570 |
| TMEM158 rabbit anti-human/mouse | Antibodies-Online | ABIN1386582 |
| **Antibodies used for Western blotting** | | |
| *Antigen* | *Vendor* | *Catalogue number* |
| STAT5 clone D2O6Y rabbit anti-mouse | Cell Signaling | 94205 |
| Phospho-STAT5 (Tyr694) clone D47E7 rabbit anti-mouse | Cell Signaling | 4322 |
| **Neutralizing antibodies** | | |
| *Antigen* | *Vendor* | *Catalogue number* |
| Csf2r rat anti mouse | RnD Systems | MAB6130 |
| **Antibodies used for flow cytometry** | | |
| *Antigen* | *Vendor* | *Catalogue number* |
| CD45 clone 30-F11 fluorophore A700 | BD Biosciences | 103128 |
| CD11b clone M1/70 fluorophore BV605 | BD Biosciences | 563015 |
| Ly6G clone 1A8 fluorophore BV421 | BD Biosciences | 562737 |
| Ly6C clone HK1.4 fluorophore PerCP Cy5.5 | Biolegend | 128011 |
| CD49d clone R1-2 fluorophore Pe-Cy7 | Biolegend | 103618 |
| CD4 GK1.5 fluorophore PE-Vio770 | Miltenyi | 130-102-784 |
| CD8 clone 53-6.7 fluorophore PerCP-Cy5.5 | BD Biosciences | 561109 |
| Asca2 clone REA969 fluorophore APC | Miltenyi | 130-117-535 |
